# Unravelling ATP processing by the AAA+ protein p97 at the atomic level

**DOI:** 10.1101/2023.08.26.554927

**Authors:** Mikhail Shein, Manuel Hitzenberger, Tat Cheung Cheng, Smruti R. Rout, Kira D. Leitl, Yusuke Sato, Martin Zacharias, Eri Sakata, Anne K. Schütz

**Affiliations:** Faculty for Chemistry and Pharmacy, Ludwig Maximilian University of Munich, 81377 München, Germany; Bavarian NMR Center, Technical University of Munich, 85748 Garching, Germany; Institute of Structural Biology, Helmholtz Zentrum München, 85764, Neuherberg, Germany; Physics Department and Center of Protein Assemblies, Technical University of Munich, 85748 Garching, Germany; Institute for Neuropathology, University Medical Center Göttingen, 37077 Göttingen, Germany; Multiscale Bioimaging: from Molecular Machines to Networks of Excitable Cells (MBExC), University of Göttingen, 37073 Göttingen, Germany; Institute for Auditory Neuroscience, University Medical Center Göttingen, 37077 Göttingen, Germany; Center for Research on Green Sustainable Chemistry, Graduate School of Engineering, Tottori University, Tottori, Japan; Department of Chemistry and Biotechnology, Graduate School of Engineering, Tottori University, Tottori, Japan

## Abstract

The human enzyme p97 regulates various cellular pathways by unfolding hundreds of protein substrates in an ATP-dependent manner, making it an essential component of homeostasis and impactful pharmacological target.

The hexameric complex undergoes substantial conformational changes in the course of its catalytic cycle. Here, we elucidate the molecular motions that occur at the active site in the temporal window immediately before and after ATP hydrolysis by merging cryo-EM, NMR spectroscopy and MD simulations. p97 populates a metastable reaction intermediate, the ADP.Pi state, which is poised between hydrolysis and product release. Detailed snapshots reveal that the active site is finely tuned to trap and eventually discharge the cleaved phosphate. Signalling pathways originating at the active site coordinate the action of the hexamer subunits and couple hydrolysis with allosteric conformational changes.

Our multidisciplinary approach enables a glimpse into the sophisticated spatial and temporal orchestration of ATP handling by a prototype AAA+ protein.

---

The ATP-dependent enzyme p97 powers diverse energy-consuming processes in the cell. p97 is a homo-hexamer in which each subunit comprises two ATPase domains, D1 and D2, that assemble into two stacked rings (Fig. 1a). Its N-terminal domain (NTD) recruits cofactors and substrates and is positioned according to the nucleotide bound in D1: elevated above the D1-ring when ATP is bound (NTD ‘up’) and coplanar in ADP-bound form (NTD ‘down’)^1, 2^. As a result, the NTD undergoes a large-scale motion during the ATP-hydrolysis cycle. p97 is member of the ATPases associated with diverse cellular activities (AAA+) superfamily that features conserved functional elements for nucleotide binding and hydrolysis such as the Walker A and B motifs, the arginine-finger and the sensor motifs^3^. As the p97 hexamer assembles, twelve active sites emerge at the inter-subunit interfaces, allowing for allosteric coordination of enzymatic activity among the subunits^3^.

**Fig. 1:**
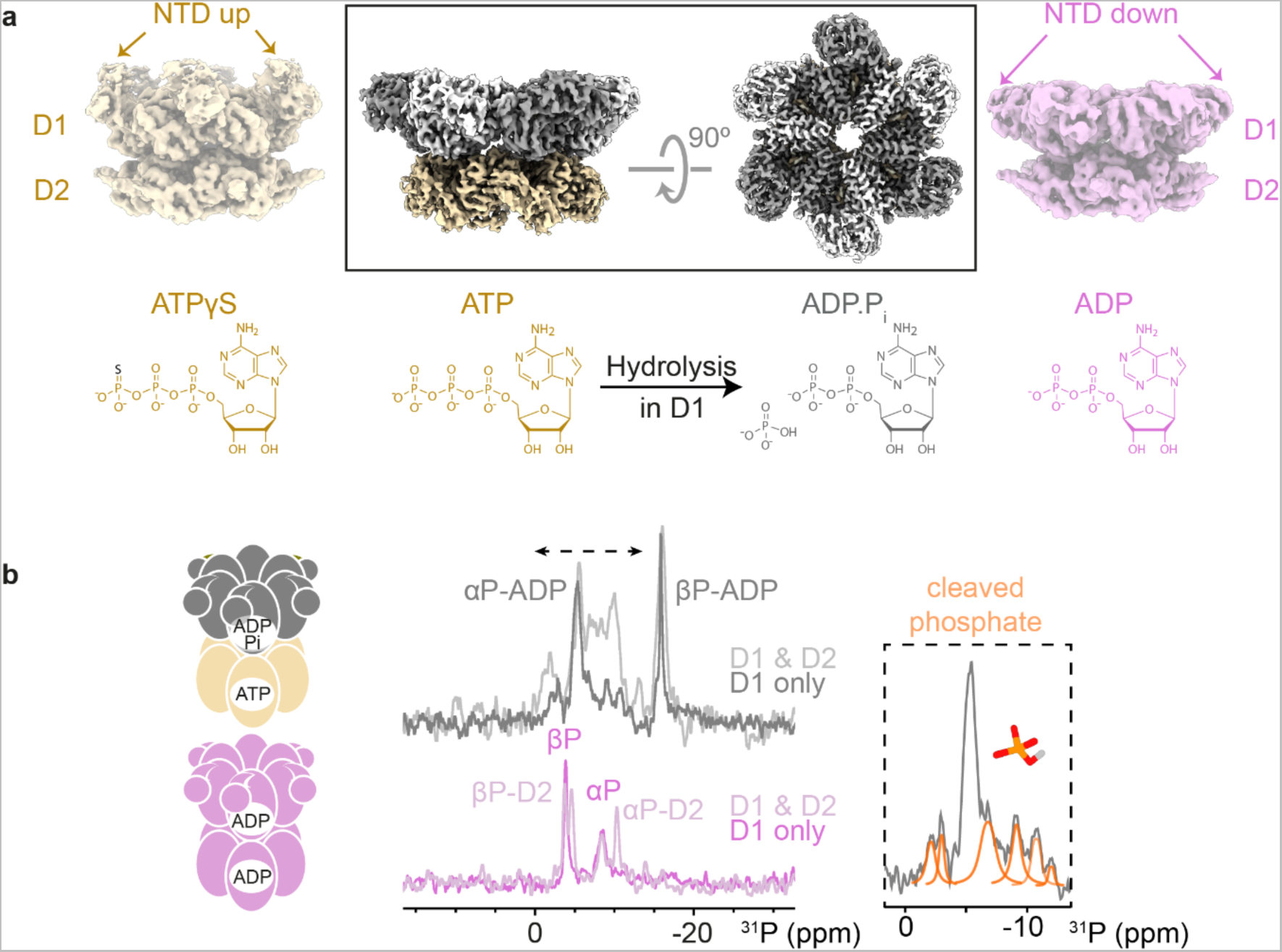
Global conformational changes linked to nucleotide turnover in the tandem ATPase p97. **a**, Box: Single-particle cryo-EM reconstruction of p97 in ADP.Pi state reveals a symmetric hexamer with NTD domains in the ‘down’ position (4 mM Mg^2+^, 10 mM ATP with regenerating system; EMDB 16781/16782, this work). For comparison, reconstructions of ATPγS- and ADP-bound p97 are shown (EMDB 3298, 3299^1^). The colouring of the cryo-EM maps reflects the bound nucleotides as shown below. Details on reconstruction in Fig. S1-3. **b,** Magic-angle spinning ^1^H→^31^P cross polarization NMR spectra of p97-bound nucleotide in the presence of ATP (top) and ADP (bottom). The p97-ND1L hexamer (residues 1-480) is ATPase active^2^ and contains only signals from D1; the corresponding spectra of fl p97 are shown in lighter hues. The observation of multiple weaker signals (orange fit) is ascribed to phosphate ions in chemically distinct environments. These signals must derive from the cleaved 𝛾-phosphate of ATP locked in the D1 active site since thio-substitution at this position in ATP results in a strong downfield shift^4^.

We previously reported that in the presence of ATP and absence of cofactors and substrates, p97 populates a uniform nucleotide state: D1 is occupied with ADP and still hosts the cleaved phosphate (P_i_) ion^4^. Conformational analysis by nuclear magnetic resonance spectroscopy (NMR) indicated that the observed state is distinct from apo, ADP- or slowly-hydrolysable ATP𝛾S states. A reaction intermediate in which the bond between 𝛾- and β-phosphate groups of ATP has been cleaved but neither reaction product released was postulated 50 years ago and termed ‘ADP.P_i_‘ state^5^. Due to its limited lifetime, it has been little characterized at the atomic level.

Single-particle cryo-electron microscopy (cryo-EM) enables structural analysis of such short-lived species. While resolutions below 4 Å are sufficient to establish the identity of the nucleotide ^1, 6–8^, *i.e.* whether ATP or ADP is bound, it remains challenging to establish the location of the cleaved P_i_ ion as its distance to the nucleotide is not known *a priori* and its location may fluctuate, causing smearing of the cryo-EM density. Exceptions are Hsp70^9^, F-actin^10^, myosin^11^ and F_1_-ATPase^12^, which all form stable ADP.P_i_ adducts with exogeneous P_i_ ions that may not reflect the authentic reaction intermediates preceded by enzymatic hydrolysis events. So far, no transient ADP.P_i_ structure after P_i_ cleavage has been reported or recognized as such.

Here, we derive the structure of ADP.P_i_-bound p97 via cryo-EM and molecular dynamics simulations (MD). This snapshot of ATP processing reveals how the active site first accommodates and then releases the cleaved P_i_ ion. We dissect the contributions of active site residues and identify the underlying triggers that induce domain motion upon hydrolysis. Additionally, we map pathways that coordinate activity between adjacent subunits. Our investigation sheds light on the structural transitions and dynamical changes that accompany ATP processing by multimeric enzymes.

## Results

### Observation of a post-ATP-hydrolysis reaction intermediate

Full-length (fl) p97 at physiological Mg^2+^ ion and ATP concentrations in the presence of an ATP-regeneration system was flash frozen and subjected to single-particle cryo-EM. The reconstructed hexamers display no deviation from sixfold symmetry with all NTDs positioned coplanar with the D1 ring. A C6-symmetrized reconstruction pushed the final resolution to 2.61 Å (Fig. 1a). Overall, the structure of the D1 domain is similar to that of ADP-bound p97^1^. Elements related to NTD ‘down’ state are fully built, notably the helix-loop conversion in the NTD-D1 linker and NTD-D1 interfaces. The nucleotides in D1 and D2 were assigned to ADP and ATP, respectively (Fig. E1). While the ATP molecule is clearly defined, weak cryo-EM densities are observed around the ADP molecule in D1, hinting at the presence of additional molecules and structural heterogeneity.

To confirm the identity of the p97-bound nucleotide, we acquired ^31^P NMR spectra of nucleotide bound to p97 during ATP turnover (Fig. 1b). Comparing the spectra acquired on fl p97 to a variant lacking D2 domain (p97-ND1L, residues 1-480^2^), the ⍺-phosphate (P) and β-P signals of the ADP molecule in D1 can be assigned. In addition, multiple weaker signals are attributed to P_i_ ions trapped at the active site in a heterogenous environment. EM and NMR concur that a metastable ADP.P_i_ nucleotide state has been captured in D1, which we subjected to in-depth structural analysis.

### Structure of the active site in ADP.P_i_ state

Unlike the clearly defined phosphate moieties of the ATP molecule in D2, the cryo-EM density of ADP in D1 is tailed, with unexplained densities emanating from the β-P (Fig. 2a, E1) and close to the arginine finger R359. We obtained a trajectory of the P_i_ and Mg^2+^ ions immediately after ATP hydrolysis from MD simulations. Starting from ATP-bound p97 hexamer, the ATP molecule in one of the six subunits was converted to ADP.P_i_ *in silico* followed by 2 µs of unrestrained simulation. After rearrangements at the active site within the first few nanoseconds, two clusters emerge indicating stable positions of the Mg^2+^ and P_i_ ions: *(i)* In the first cluster, termed state A, the leaving P_i_ ion is stabilized by Walker A residue K251 as well as sensor residue N348. R359 binds to P_i_ but sometimes dissociates or binds via water. It is much more mobile than N348, which maintains a persistent binding mode with respect to the P_i_ ion. *(ii)* In the second cluster, termed state B, the leaving P_i_ is detached from K251 and positioned closer to R359 and R362, thus being pulled towards the adjacent, *trans*-acting subunit. The clusters superimpose well with the unassigned cryo-EM densities (Fig. 2a).

**Fig. 2:**
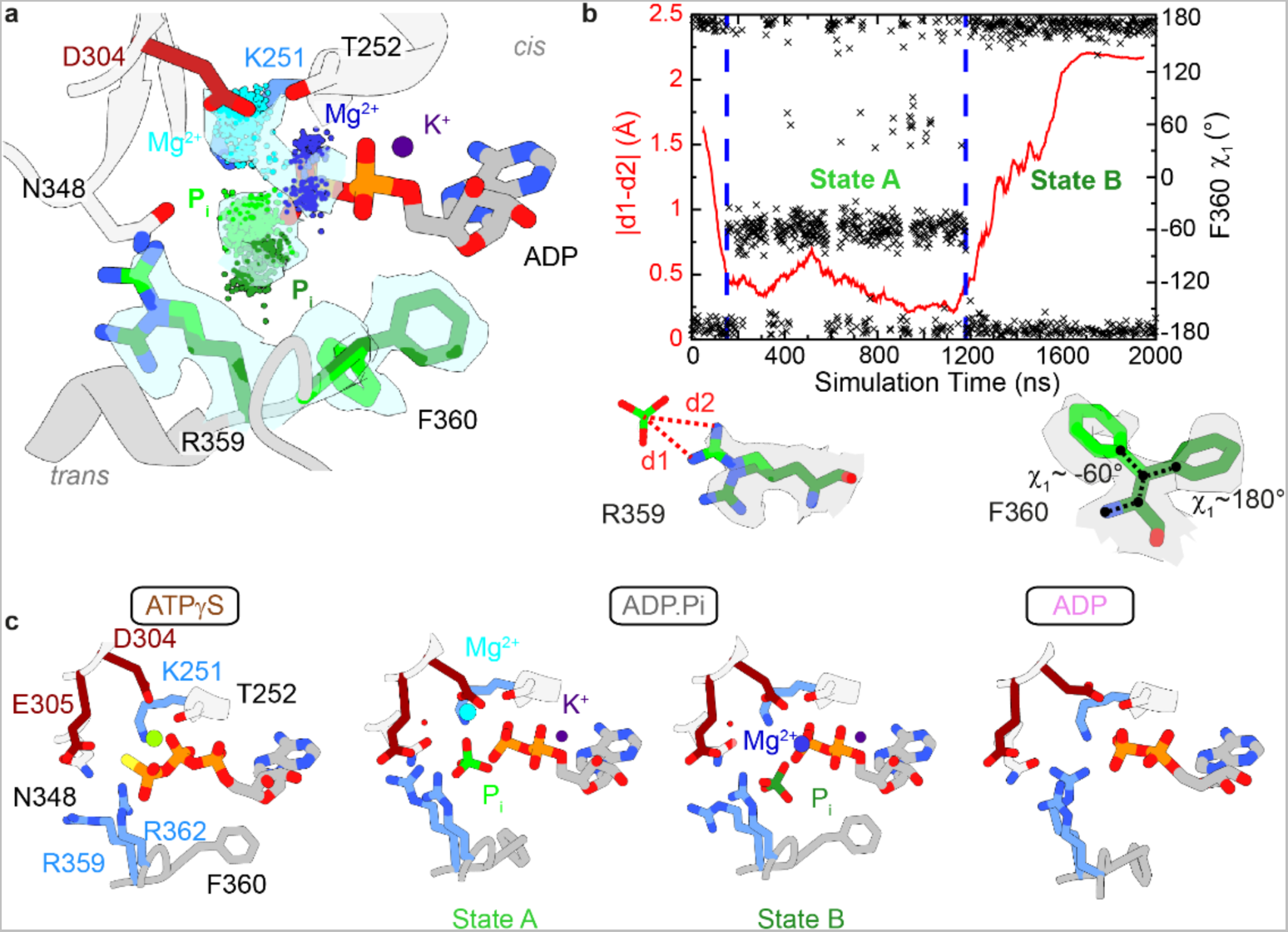
Coordination of the cleaved Pi ion in D1. **a**, Zoom on the unaccounted density at the D1 active site. Snapshots from the MD trajectory evaluated at 2 ns intervals identify at least two locations each for Pi and Mg^2+^. The convergence between MD and cryo-EM enables the assignment of Mg^2+^ (cyan for state A, dark blue for state B), cleaved Pi and R359/F360 rotamers (light for state A, dark green for state B). The iterative modelling process is outlined in Fig. E3. **b**, In MD simulations of the ADP.Pi state, R359 and F360 undergo a correlated motion on a µs-time scale, evidenced by fluctuations of the side chain dihedral angle (F360 ξ1) and the phosphate-arginine binding geometry. A transition between the two stable geometries, states A and B, occurs here after ∼ 1200 ns. The side chain rotamers are visible in the experimental cryo-EM density. **c**, Juxtaposition of the D1 nucleotide binding pocket in ATPγS (PDB: 5ftn^1^), ADP (PDB: 5ftk^1^) and ADP.Pi states (PDB: 8cpf, this work).

*In-silico* analysis suggests that the Mg^2+^ ion stabilizes the leaving P_i_, compensating the Coulombic repulsion from the β-P of ADP. In both states, an octahedral coordination geometry of the Mg^2+^ ion is achieved (Fig. E2a,b). Compared to the ATP state (Fig. E2c), the Mg^2+^ ion dissociates from T252 and instead interacts with Walker B residue D304, either directly or bridged via water molecules. With regard to the protonation state of the leaving P_i_ ion, only the simulation featuring HPO_4_^2−^ is in agreement with the experimental cryo-EM density of the ADP.P_i_ state, while the simulation featuring H_2_PO_4_^−^ exhibits conformations and dynamics nearly identical to those of the ATP state. (Fig. E2d,e).

In our cryo-EM map of the ADP.P_i_ state, the side chains of three residues at the active site exist in two rotamers: R359 and F360 that interact with the nucleotide in *trans* as well as H384 in *cis* (discussed below). In the simulations, the F360 rotamer motion is correlated to the interaction mode between the cleaved phosphate and R359 (Fig. 2b, Video S1). The head-on bidentate complex of the P_i_ ion with two amino groups in state A is linked to the F360 χ_1_ = 180° conformer while the lateral monodentate complex of R359 in state B is linked to the χ_1_ = -60° conformer.

By iterative integration of MD and EM, we determined the positions of the leaving P_i_ and Mg^2+^ ions as well as the associated conformations of active-site residues (Fig. E3). The following features set apart the ADP.P_i_ state from the ADP and ATP𝛾S states (Fig. 2c): the active site is heterogeneous with at least two distinct positions for P_i_ and Mg^2+^ ions; K251 interacts more with the leaving P_i_ than with ADP; the Mg^2+^ ion has dissociated from T252 to interact with D304; N348 coordinates the P_i_ ion; R359 and F360 occupy two side chain rotamer states, reflected in µs-time-scale motion in MD simulations.

**Fig. 3:**
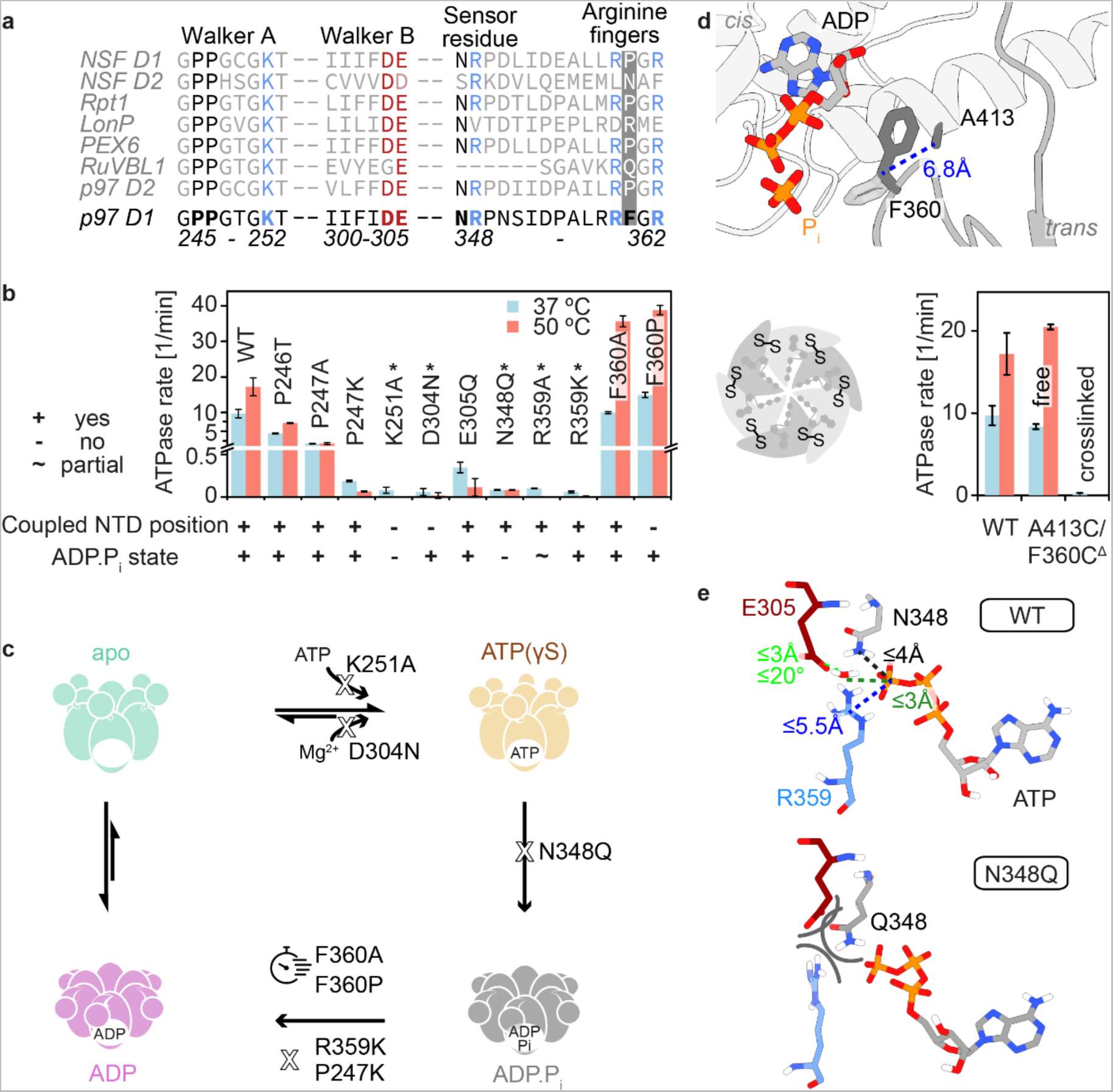
Function of active-site residues in the ATP-hydrolysis cycle. **a**, Sequence alignment of human AAA+ family proteins. While several key motifs are highly conserved, F360 is unique to the p97 D1 domain (Fig. S4). **b**, ATPase rates of p97-ND1L bearing point mutations at the active site and their functional defects deduced from NMR analysis. All presented mutants form hexamers; only N348Q fully abolishes ATP hydrolysis; only mutations of F360 have a stimulatory effect on phosphate release; *designates ATPase inactive mutants. Coupled NTD position’ indicates whether the mutant exhibits the same change in NTD position upon nucleotide binding as wild type (wt) p97. ‘ADP.Pi state’ indicates that this state is observed during ATP turnover. **c**, Impact of mutations on the four steps of the ATP-hydrolysis cycle. **d**, In ADP.Pi state, F360 from the *trans*-acting subunit samples two side chain rotamer states, one of which contacts helix ⍺407-423 of the active subunit. Crosslinking of C360 to this helix at C413, but not the mutations alone, abolishes ATPase activity of D1. ^Δ^ designates a cysteine-free p97 variant. **e**, Criteria that define hydrolysis-active conformations^16^, amended for p97: *i*) A water molecule next to the terminal phosphate (dark green) forms a hydrogen bond to E305 side chain (light green). This lytic water molecule is polarized and thus activated for attack. *ii*) R359 polarizes the 𝛾-phosphate and is poised to hydrogen bond after cleavage (blue). *iii*) The 𝛾-phosphate is held in place by N348 via a hydrogen bond (black). Simulations of the N348Q mutant lack hydrolysis-active conformations due to steric hindrance from the longer Q side chain.

### Contribution of individual residues to the processing of ATP

The D1 domain contains both signature AAA+ motifs and elements unique to p97 (Fig. 3a). To explore the roles of the active-site residues, we conducted biophysical assays on point-mutated p97-ND1L. Each mutant was subjected to a stepwise assessment of defects in assembly, nucleotide binding, and ATPase activity (Fig. E4). We further mapped the conformational dynamics as a function of bound nucleotide by NMR (Fig. E5).

**Fig. 4:**
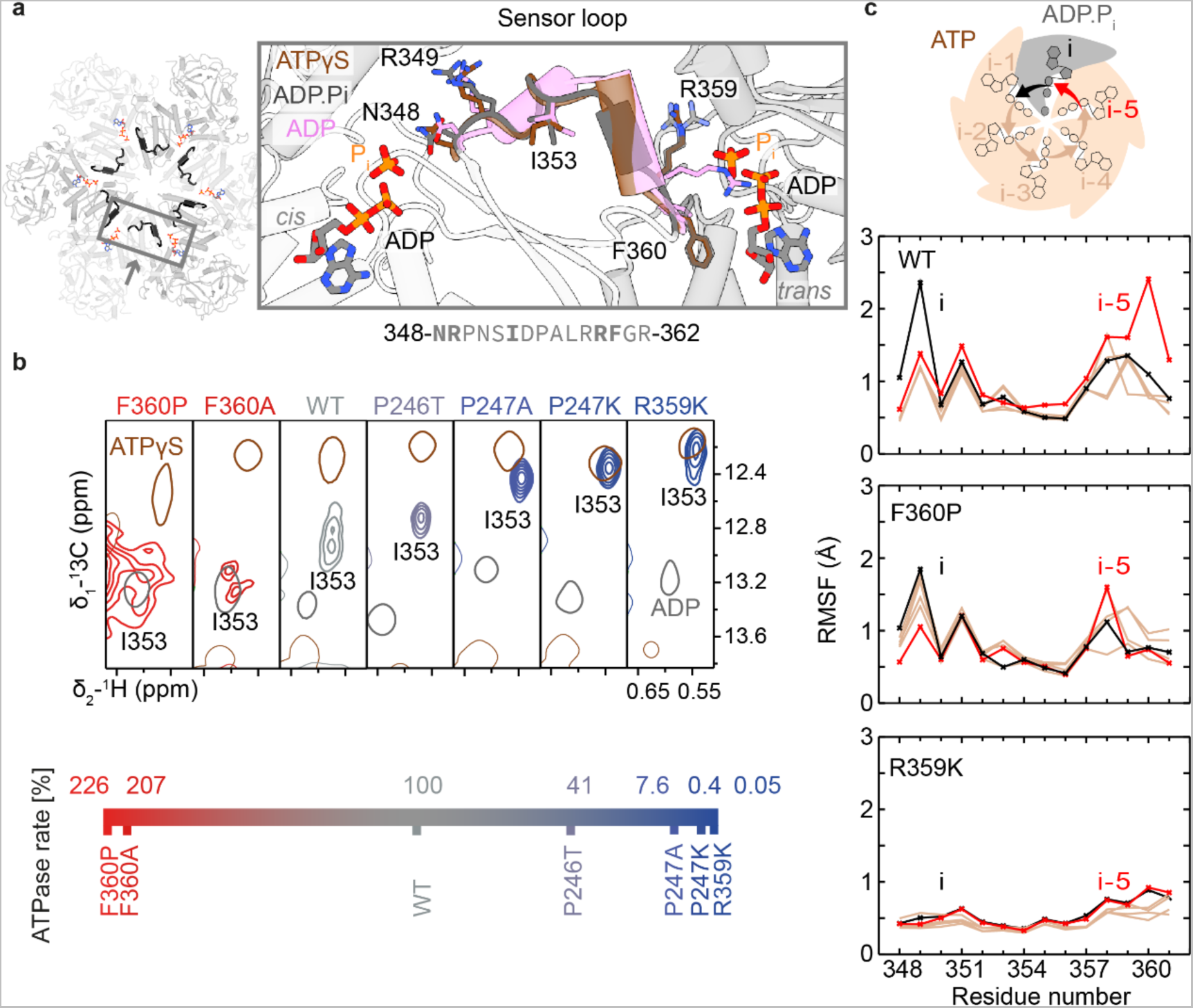
Inter-subunit communication channel connects active sites. **a**, The sensor loop bridging the N348 of one nucleotide binding pocket to F360 of the anticlockwise adjacent pocket changes conformation between ATP(𝛾S) and ADP-bound states. In the ADP state, R349-I353 form a 310 helix; in the ADP.Pi state, this transition is incomplete, making this loop the last structural element to convert after ATP hydrolysis (Ramachandran analysis in Fig. E7) **b**, The signal of the I353 Cδ1-methyl group in the middle of the loop displays line broadening in the ADP.Pi state of wt p97. For hydrolysis-competent mutants, the ATPase activity correlates with the extent of conversion from an ATP𝛾S-like to an ADP-like conformation. This correlation could reflect the coupling of loop motions to product release, the rate-limiting step of the ATP-hydrolysis cycle. **c**, C⍺-RMSF fluctuations quantify the deviation of residues from their average position over the course of the 2 µs MD trajectory. In wt p97, mobility is pronounced in sensor loops neighbouring ADP.Pi-bound but not ATP-bound pockets. In the hyperactive F360P mutant, mobility is increased in all subunits, irrespective of nucleotide state; in the hypoactive R359K mutant it is strongly decreased in all subunits. The corresponding RMSD analysis is shown in Fig. S10, excerpts from the MD simulations in Videos S2, S3.

**Fig. 5:**
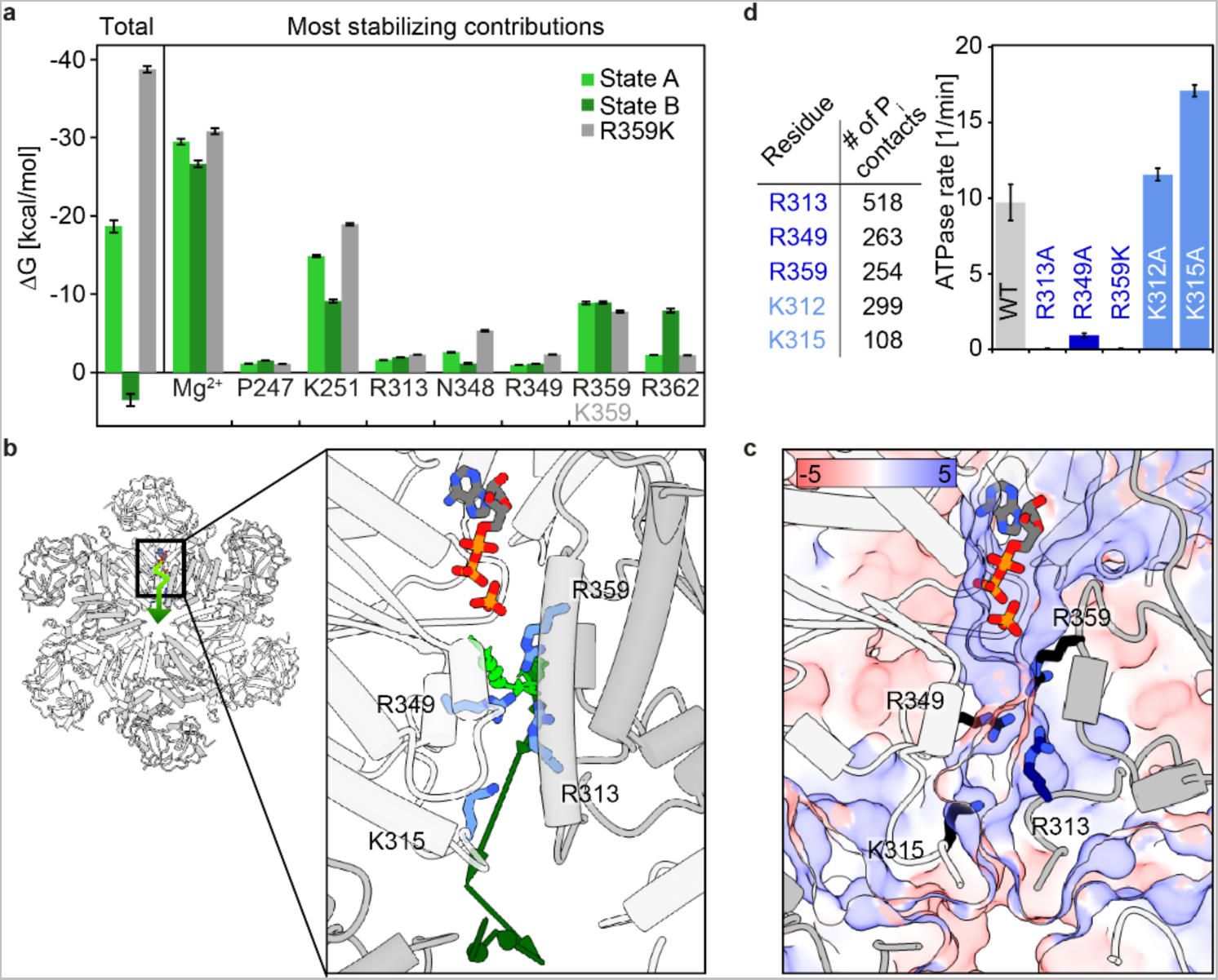
Pathway of phosphate dissociation. **a**, Free energies of ADP.Pi complexes from MMPBSA^20^ calculations. Energy decomposition identifies entities that stabilize the leaving Pi ion most strongly: the Mg^2+^ ion and the side chain of K251. Electrostatic terms but not solvation terms are responsible for energy differences between the states. **b**, Illustration of phosphate dissociation (arrows, light to dark green) from the active site towards the central pore derived from an MD trajectory; side view in Fig. E8c. **c**, Same view as panel b, colour coded according to electrostatic potential in units kT/e (APBS^21^), defines a positively lined channel. **d**, Left: positively charged residues in D1 with the highest number of contacts to the dissociating Pi (defined by a distance below 3 Å in a single frame) in the MD trajectory. Right: ATPase activities of the corresponding p97-ND1L point mutants. The mutation of R but not K residues causes a drastic decrease in ATPase activity (additional mutants in Fig. E8d.)

The results are summarized in Fig. 3b,c. Briefly, all mutants except K251A bind ADP and ATP𝛾S; all mutations except F360A/P reduce the ATPase activity. The NTD position (‘up’ vs. ‘down’) is linked to nucleotide state (apo/ATP *vs.* ADP) with the exception of D304N and F360P, which assume the ‘down’ conformation in the presence of slowly hydrolysable ATP analogues. The ‘up’ conformation of the apo state is not compromised in any mutant. Before ATP hydrolysis, the side chain of D304 hydrogen bonds with water molecules coordinating the Mg^2+^ ion. Removing its charge leads to loss of Mg^2+^ and concomitant failure to recognize bound ATP and assume the ‘up’ conformation (Fig. S6).

In the ADP.P_i_ state, F360 equally populates two rotamers while the static ATPγS state shows a preferential F360 χ_1_ dihedral of 180^1, 13^. This is echoed in the MD simulations, where it is only upon hydrolysis that F360 is unlocked and transiently dissociates from the helix ⍺_407-423_ (Fig. E6). This ability of F360 to pull the arginine finger loop towards helix ⍺_407-423_ could be essential to lock the NTD into the ‘up’ state. The critical role of F360 is underpinned by its conservation in p97 homologues but absence in AAA+ proteins without an NTD (Fig. 3a, S4). Disease-associated p97 mutants lack this rotamer switch^14^, display a dynamic NTD^15^ and no long-lived ADP.P_i_ state^4^. F360 is the only site where mutation entails a gain of ATPase function. Mobility at this site is indeed linked to ATP processing: crosslinking C360 to C410 abolishes ATPase activity (Fig. 3d).

**Fig. 6:**
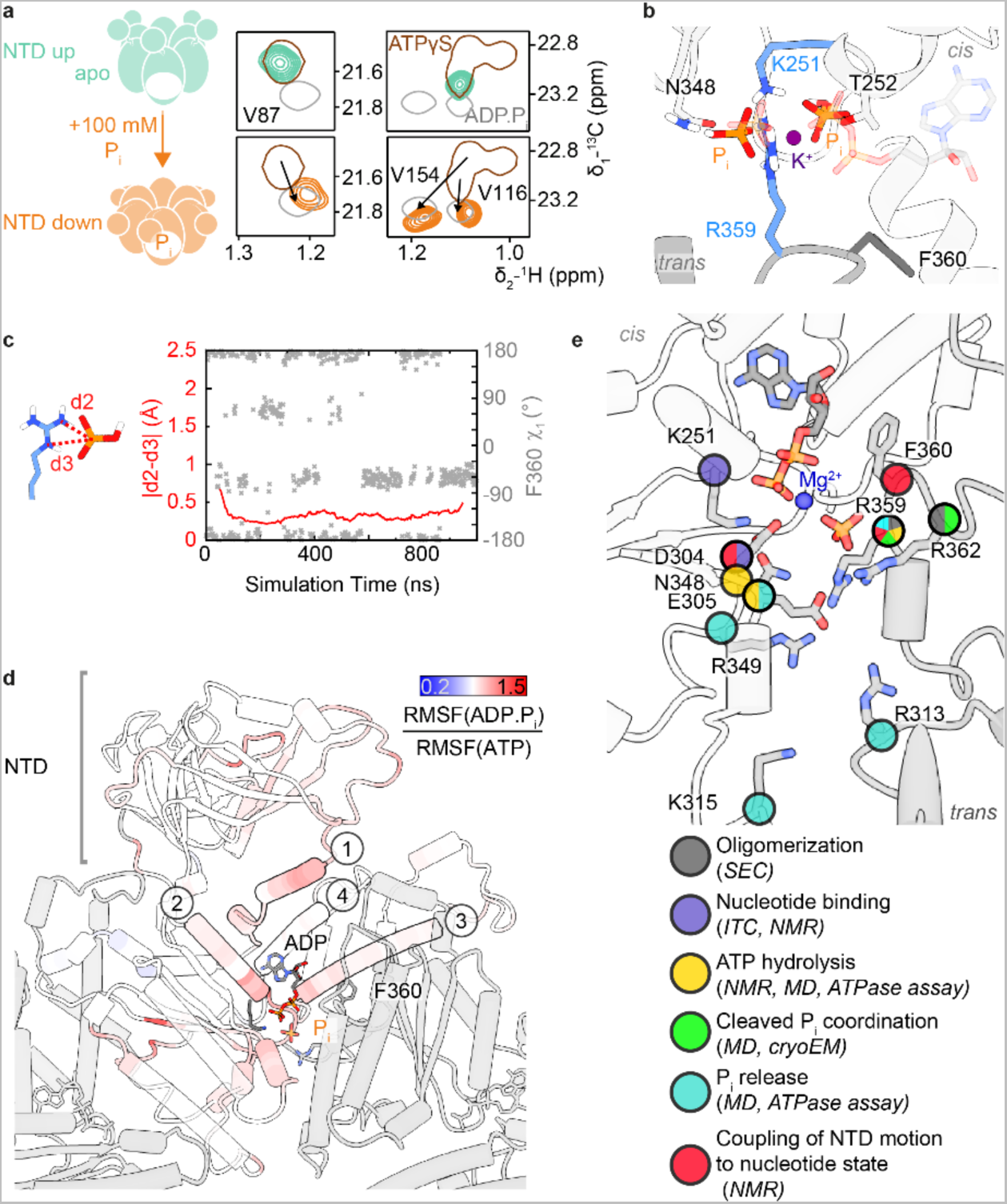
Allosteric control of NTD motion. **a**, NMR probes indicate a conformational change of the NTD induced by addition of 100 mM Pi ions to apo p97, reflected by peak shifts from ‘up’ (ATP𝛾S-like) to ‘down’ (ADP.Pi-like) position (titration in Fig. E9a). This Pi concentration is above physiological intracellular levels. The same effect was observed for arsenate and sulphate ions (Fig. E9b). **b**, Snapshot from MD simulations (Video S4) of the apo D1 nucleotide binding pocket. Two Pi ions mimic the ADP.Pi state (transparent) and occupy the same positions as the β-P of ADP and the leaving Pi, bridged by a K^+^ or Na^+^ ion from the solvent. No Mg^2+^ ion is necessary to stabilize this arrangement in simulation or experiment. **c**, The MD trajectory of Pi-bound p97 reveals a mobile F360 side chain that switches between rotamers corresponding to states A and B (Fig. 2b); R359 stably coordinates a Pi ion via its Nε atom and one amino group. **d**, Top view of one D1 subunit colour coded according to the ratio of backbone RMSF of ADP.Pi state B over ATP state, each sampled over 800 ns. Red colour indicates an increase in mobility upon hydrolysis, observed in (1) helix ⍺191-199 and the NTD-D1 linker, (2) helix ⍺251-262 extending from the Walker A motif to the NTD-D1 interface, (3) helix ⍺407-423 to which F360 associates transiently, (4) helix α374-387 running past the adenine moiety of the nucleotide (increased mobility in state A only, *cf*. Fig. E10). **e**, Summary of the function of residues at the p97 D1 active site. Categories were assigned either according to structural contributions evident from MD or cryo-EM or according to mutagenesis-induced defects in ATP processing.

### Determinants of ATP-hydrolysis competence

Real-time NMR establishes that mutants with low ATPase activity (P246T, P247A/K, E305Q, R359K, Fig. 3b) still form an ADP.P_i_ state, pointing to slow product release but intact ATP hydrolysis. However, the N348Q mutant with no measurable ATPase activity displays only the NTD ‘up’ state in the presence of ATP (Fig. S8). The sensor residue N348 is thought to position the water molecule for nucleophilic attack on ATP^17^. To recapitulate the suppression of ATP hydrolysis, we evaluated the frequency of reactive conformations at the D1 active site in MD simulations of wild type (wt) *vs.* N348Q p97 (Fig. 3e). While three of five ATP-bound subunits sampled reactive conformations with high frequency in the wt, all but one subunit were practically inactive in the mutant (statistics in Fig. S9). The longer side chain of Q348, which congests the active site, disfavors the proper geometry for ATP hydrolysis. E305 is thought to activate a water molecule for attack on the 𝛾-phosphate of bound ATP^16, 17^. The E305Q mutation strongly reduces the ATPase activity of D1^14^; still, the rate-limiting step of the catalytic cycle of this mutant remains product release^4^.

### Dynamics in the sensor loop are coupled with product release

Sequential ATP hydrolysis around the multimer ring has emerged as a plausible operation mode for AAA+ proteins^3, 18^. A communication line between the active sites of adjacent subunits must underlie such coordination. We hypothesized that the ‘sensor loop’ (Fig. 4a) could assume this function in p97 D1. Part of this loop converts from turn to 3_10_-helix between ATP𝛾S and ADP states. The ADP.P_i_ state, however, still exhibits a conformation similar to ATP𝛾S, unlike the rest of the D1 domain (Fig. E7). Transitions of the loop can be monitored via the central reporter residue I353. Its NMR signals are distinct in ATP𝛾S and ADP states and exchange-broadened in ADP.P_i_ state (Fig. 4b), indicative of a loop motion occurring on a millisecond time scale. A mutant series reveals a correlation between the extent of turn-helix conversion and the ATP-turnover rate. Globally, all mutants display the spectral signature of the ADP.P_i_ state with NTD in ‘down’ conformation. The I353 signals of the hyperactive F360P/A mutants are significantly broadened; at the other extreme, the signal of the hypoactive R359K mutant overlaps with the ATP𝛾S state. Apparently, the loop does not respond to ATP hydrolysis with a structural or dynamical change in this mutant.

We compared the residue-wise C⍺ root mean square fluctuation (RMSF) of the sensor loop in the MD simulation of a p97-ND1L hexamer with five ATP and one ADP.P_i_ bound subunits (Fig. 4c). For the wt, ATP hydrolysis increases structural fluctuations: the two subunits lining the ADP.P_i_ active site display distinct profiles with increased mobility. In simulations of mutant p97, however, loop mobility is increased for the hyperactive F360P and decreased for the hypoactive R359K mutant irrespective of nucleotide state. The RMSF of R349 in the wt significantly increases when the adjacent active site is in ADP.Pi state. The cryo-EM densities of R349 in the ADP.P_i_ map are not well defined, suggesting residual flexibility.

In sum, NMR and MD analyses concur that mobility and propensity for 3_10_-helix formation in the sensor loop are linked to the ability of product release. As the loop directly connects adjacent active sites, it is conceivable that its structural transition triggers sequential ATP hydrolysis events, which have been observed when p97 is working asymmetrically in the presence of cofactors and substrates^19^.

### Energetics of phosphate binding and release

The ADP.P_i_ state of the R359K mutant is particularly long-lived, suggesting that a specific interaction mode of the P_i_ ion with R359 might be a prerequisite to induce product dissociation. In MD simulations, its guanidinium moiety interacts exclusively with the P_i_ ion and not with ADP. In contrast, the side chain of K359 preferentially coordinates between the P_i_ and the β-P of ADP, where it shields negative charges and stabilizes the ADP.P_i_ complex in a similar manner as the Mg^2+^ ion (Fig. E8b, Video S2). While the K359 mutant populates only state A, the wt features transitions between states A and B.

To quantify the stability of wt states A *vs*. B *vs*. K359 mutant, we conducted MMPBSA (Molecular Mechanics Poisson-Boltzmann Surface Area^20^) calculations to estimate the interaction energy of the P_i_ ion to the ADP.P_i_ state of p97. This method also allows decomposition of total free energies into the most stabilizing (Mg^2+^, K251; Fig. 5a) and destabilizing (ADP; Fig. E8a) contributions. We here consider the P_i_ ion as the ligand and p97-ADP-Mg^2+^ as the receptor. Although R359 interacts with the P_i_ ion both in the simulations and the cryo-EM, it is not necessary for achieving a stable ADP.P_i_ state; the presence of a Mg^2+^ ion bridging ADP and P_i_ is sufficient. In contrast to the stable binding pose in state A of wt (ΔG ∼ - 19 kcal/mol) and K359 mutant (ΔG ∼ -39 kcal/mol), P_i_ binding is predicted as unstable in state B (ΔG ∼ +3.5 kcal/mol), chiefly due to the repulsion between ADP and P_i_. Thus, transitions from state A to B could mark the onset of P_i_ dissociation events in p97 wt. In contrast, the R359K mutant has no state B equivalent with positive ΔG and an even stronger stabilization of the P_i_ ion compared to wt state A. This combination manifests in inefficient product release and low ATP turnover rates.

Over the 2 µs-simulation, the ADP.P_i_ state remains stable as long as cations bridging ADP to P_i_ are present. We therefore expedited complex dissociation by removing the Mg^2+^ ion artificially. In the resulting trajectory (Fig. 5b), the P_i_ travels from the active site towards the centre of the hexamer to dissociate through the central pore along a channel lined by positive charges (Fig. 5c). Intriguingly, the ion is handed over from R359 of the *trans*-acting subunit to R349 of the *cis*-acting subunit, which mark the end and the start of the respective sensor loops. To probe the effective contributions of individual residues to P_i_ evacuation, we generated a series of point mutants altering charges or exchanging R↔K and determined their ATPase activities (Fig. 5d, E8d). Removal of R but not K strongly reduces ATPase rates; the replacement of K by R cannot rescue slow-release mutants. In line with the MMPBSA analysis, the sequential interaction of the P_i_ ion with R359, R349, R313 could mark the onset of P_i_ release. The active site and dissociation channel are finely evolved to ensure both the initial stabilization and the eventual evacuation of the reaction products.

### Signalling of hydrolysis-induced domain motion

The NTD position is controlled allosterically by the nucleotide state in D1: in apo and ATP(𝛾S)-bound states, it is detached from and elevated above D1 domain; in ADP.P_i_ and ADP-bound states, it moves downward to form an extensive interface. This process is not observed in our MD simulation, which captures only 2 µs immediately after hydrolysis. However, we could identify the minimal structural signal to stabilize the NTD to the coplanar position by titrating P_i_ ions to apo p97-ND1L (Fig. 6a). Simulations corroborate the experimental result, suggesting that two P_i_ ions bridged by a monovalent cation bind stably to apo-p97. They adopt the same positions as P_i_ and β-P in the ADP.P_i_ state (Fig. 6b). The resulting complex reproduces the two-pronged interaction between P_i_ and the R359 guanidinium group as well as the rotamer switches of F360 χ_1_ (Fig. 6c).

From the active site, structural changes induced by ATP hydrolysis must be relayed towards the NTD where they ultimately induce a downward motion. We evaluated the RMSF of C⍺ atoms over the trajectory and visualized their ratio between ADP.P_i_ to ATP subunits as a heat map on the MD structure in Fig. 6d and E10a. The interaction between the leaving P_i_ and R359 induces the dissociation (state A) and re-association (state B) of F360 with respect to helix ⍺_407-423_ and thereby increases the plasticity of the arginine finger loop and the entire active site. This increased plasticity propagates towards the periphery of D1: first along the NTD-D1 linker; second along the helix extending from the Walker A motif towards the NTD; third along helix

⍺_407-423_ from the ribose moiety towards the *trans*-acting subunit; fourth from the adenine moiety along helix α_374-387_ towards the NTD. The latter effect is reflected in the cryo-EM map: only in ADP.P_i_ state does H384 display a second side chain rotamer that contacts the ribose. The nucleotide is thus slightly repositioned, helix α_374-387_ shifts with respect to ATPγS state, leading to a flipping of N387 located at the end of the helix. The repositioning of N387 in turn enables the formation of an electrostatic network with NTD residues that fix the ‘down’ state (Fig. E10b, Video S5). Even though P_i_ ions are sufficient to evoke the ‘down’ state of the NTD, the nucleoside moiety still contributes to the signaling of ATP hydrolysis. The MD trajectory cannot capture the downward motion of the NTD, yet it pinpoints early dynamical changes that could eventually pave the way for this large-scale conformational transition.

## Discussion

Resolving the mechanism by which ATP hydrolysis is catalysed and the concomitant release of chemical energy is conveyed to mechanical motion is a major challenge in the field of enzymology. Structures of transiently captured intermediates allow dissection of the catalytic cycle into experimentally grounded snapshots. The ADP.P_i_ intermediate, in which the bond between 𝛾- and β-phosphate groups has been cleaved but neither the P_i_ ion nor ADP has been released yet, has been poorly characterized. Its existence was first postulated for myosin^5^, in which a stable ADP.P_i_ complex can be artificially induced with exogenous P_i_ – a property shared by myosin^11^, F-actin^10^ and Hsc70^9^. However, structures of such stable complexes may not reflect the authentic short-lived states during enzymatic hydrolysis nor do they cover any members of the most prevalent nucleotide-binding fold, the P-loop NTPases^22^.

We report here the 2.6 Å cryo-EM structure of the human ATPase p97 captured in a transient ADP.P_i_ state, which converges with MD simulations of the same state. Mutagenesis and NMR analyses identify the contributions of active-site residues to ATP turnover, summed up in Fig. 6e. The structures capture molecular motions that accompany ATP hydrolysis: The cleaved P_i_ travels together with the Mg^2+^ ion, which is stabilized by water molecules and coordination to D304. P_i_ release is coupled with rotamer exchanges in the arginine finger loop. A further rotamer exchange of H384 triggers a conformational transition that could ultimately effect the large-scale motion of the NTD. A stable Mg^2+^·P_i_ complex and water networks were observed for F-actin^23, 24^ and Hsc70^25, 26^, indicating a common mechanism whereby the Mg^2+^ ion plays a key role in P_i_ release. Indeed, the ADP.P_i_ state remains stable as long as bridging cations are present in simulations. Notably, in a GHL ATPase, the switch of a K residue near the nucleotide was proposed to trigger P_i_ release, which parallels our finding of an arginine finger rotamer switch^27^.

We identify a loop connecting the sensor I motif to the arginine finger of the counter-clockwise subunit. Its ability to transition from turn to 3_10_ helix between ATP, ADP.P_i_ and ADP states is correlated to efficient product release. p97 is a prototype member of the AAA+ superfamily, and ADP.P_i_ states are frequently invoked in mechanistic models of these ring-shaped oligomers^28–30^. A consensus has emerged that ATP hydrolysis proceeds counter-clockwise in substrate-engaged AAA+ proteins^3, 18^. ADP release disrupts the subunit interface and causes the respective subunit to move to the bottom of the spiral staircase and disengage from the substrate^28, 29^. Such models presume efficient inter-subunit communication elements, such as the sensor loop identified in p97.

p97 is the first AAA+ protein for which a transient ADP.P_i_ state has been captured to our knowledge. A high kinetic barrier to P_i_ dissociation, rendering release rate limiting, could be a speciality of p97-D1: the conserved F residue in the arginine finger of p97-D1 is replaced by P in p97-D2 and many AAA+. We show that F360 rotamer states regulate the NTD position and are coupled with P_i_ release. While ATP turnover in D1 is linked to NTD motion, D2 drives substrate translocation^19, 31^.

Our methodology delineates a general strategy to overcome resolution limits in the characterization of short-lived and heterogeneous enzymatic reaction intermediates. Single-particle cryo-EM affords the bulk of structure determination; MD simulations validate the interpretation of the map at the critical active site and introduce a time axis to connect multiple structural states; NMR assesses dynamical changes coupled with enzymatic events.

## Extended Data

**Extended Data Fig. 1.**
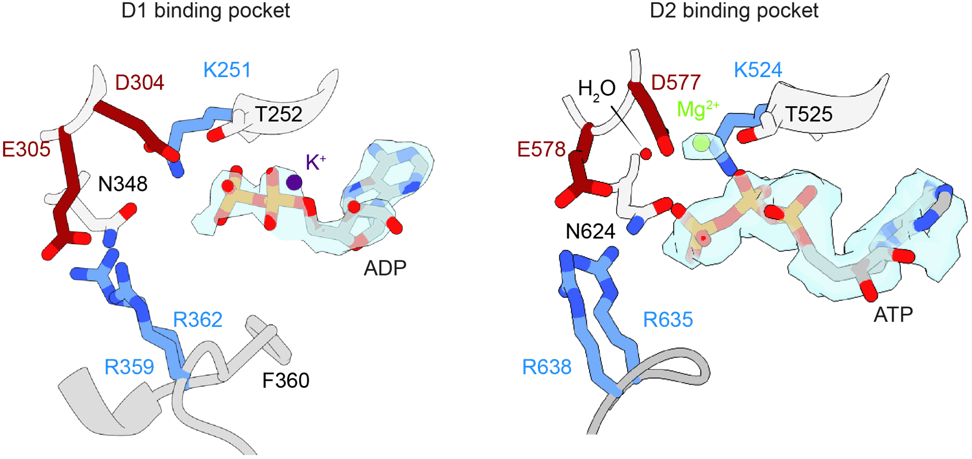
Nucleotides bound in D1 and D2. Zoom of the cryo-EM density that is assigned to the nucleotide bound to fl p97 in the presence of ATP. An ADP molecule with two phosphate groups and an ATP molecule with three phosphate groups are clearly resolved in the binding pockets of D1 and D2 domains, respectively. In MD simulations, Na^+^ or K^+^ ions are often found binding to the ⍺-P of ADP in D1, coinciding with unassigned electron density in this area.

**Extended Data Fig. 2.**
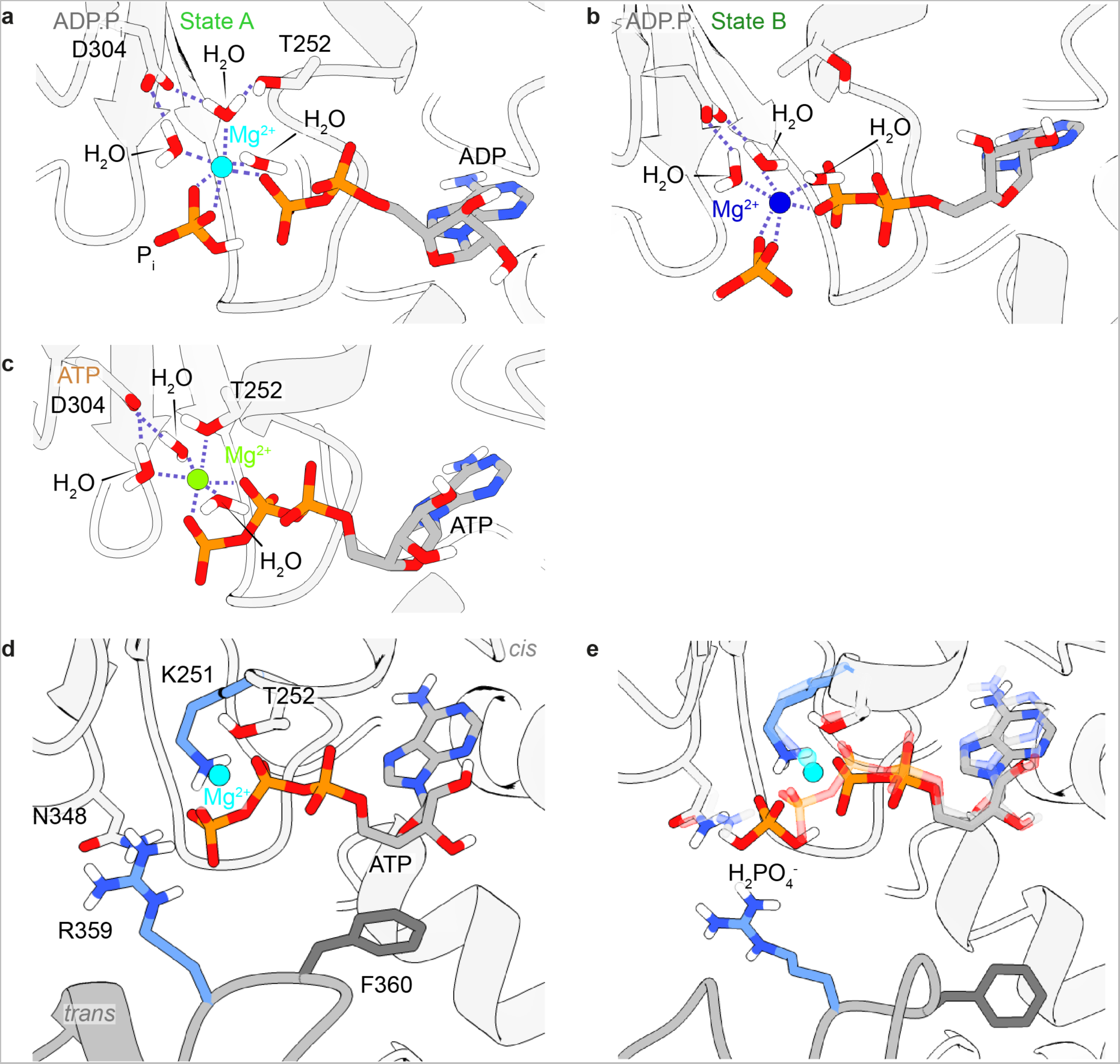
Snapshots from MD simulations of D1 in different nucleotide states. **a-c**, Octahedral coordination of the Mg^2+^ ion in ADP.Pi states A **(a)** and B **(b)** and in ATP state **(c)** in the MD simulations. **d**, The X-ray structure of ATPγS-bound p97^14^ was used as a starting structure for MD simulations with ATPγS converted to ATP, of which a snapshot is shown here. **e**, Snapshot from an MD simulation of the D1 active site containing ADP and a doubly protonated phosphate ion (H2PO4^-^) superimposed on the ATP state from panel d in transparent. The ADP.Pi state with H2PO4^-^ shows high similarity to ATP-bound state and does not coincide with the experimental electron density.

**Extended Data Fig. 3.**
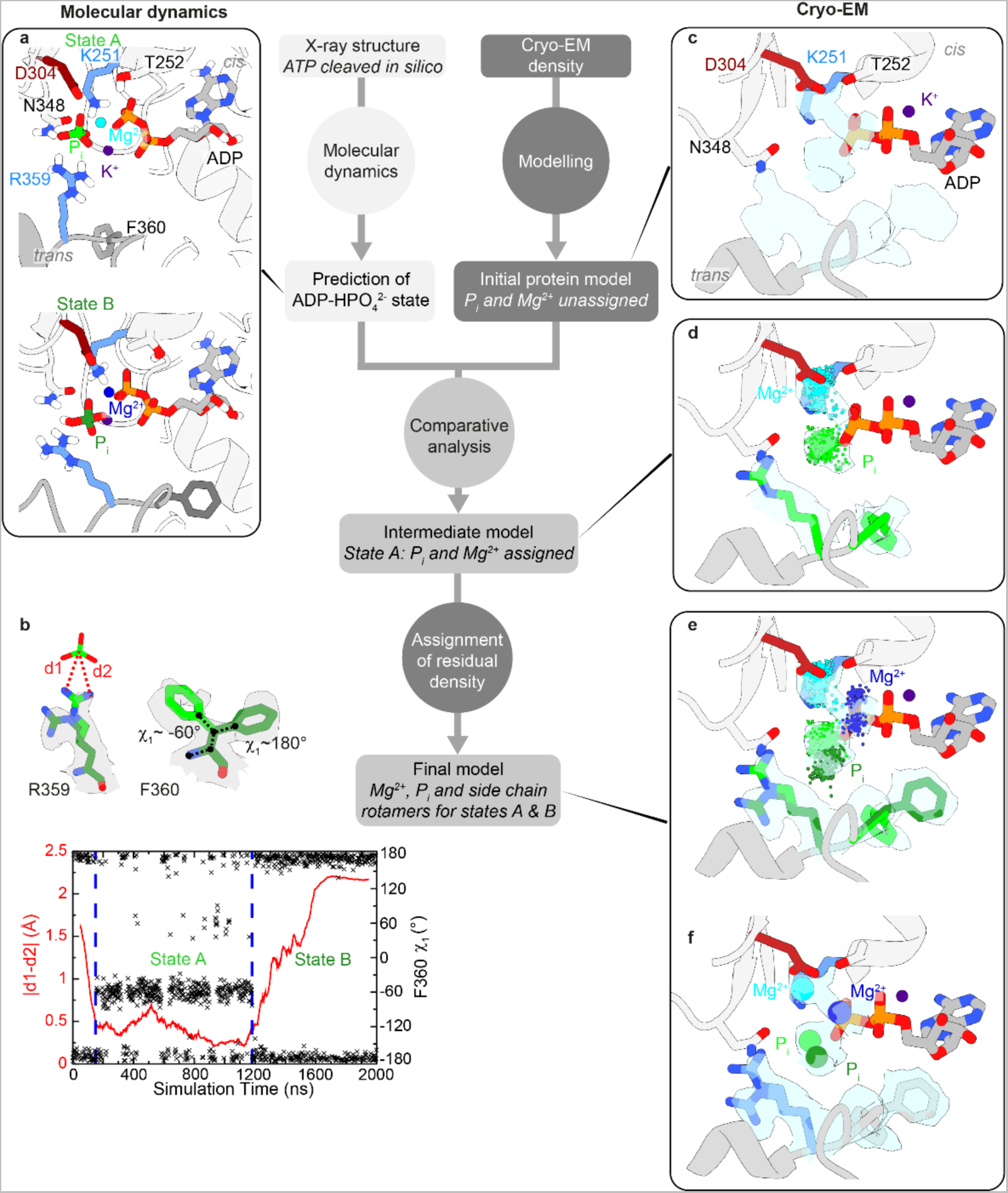
Stepwise assignment of the cryo-EM density at the D1 active site. The positions of the Mg^2+^ and Pi ions and the side chain rotamers in ADP.Pi state were determined by matching the MD simulations (left column) to the residual cryo-EM density (right column) surrounding the ADP molecule in D1. **a,** Snapshots from simulations of ADP.Pi state reveal at least two distinct stable geometries, termed states A and B, which differ in the position of the Pi and Mg^2+^ ions. **b,** The side chains of R359 and F360 undergo a correlated motion in the MD trajectory. Left ordinate: the distances d1 and d2 between R359-N^ρι^^1^/N^ρι^^2^ and the P atom of the cleaved Pi ion reflect the side chain conformation of R359. Right ordinate: ξ1 angle of F360. **c**, Residual densities at the D1 active site after assignment of the protein and ADP. Two side chain rotamers each for R359 and F360 are evident from the cryo-EM density. **d**, Snapshots taken from an MD simulation sampling state A every 2 ns superimposed on the structural model. The predicted positions of the Pi (lime) and Mg^2+^ (cyan) ions and one set of rotamers for R359 and F360 (lime) coincide with the unassigned cryo-EM density. **e**, Convergence between MD snapshots of state B and residual densities: Pi (dark green) and Mg^2+^ (blue) ions, second set of rotamers for R359 and F360 (dark green). **f,** Final structural model of the D1 binding pocket in the ADP.Pi state.

**Extended Data Fig. 4.**
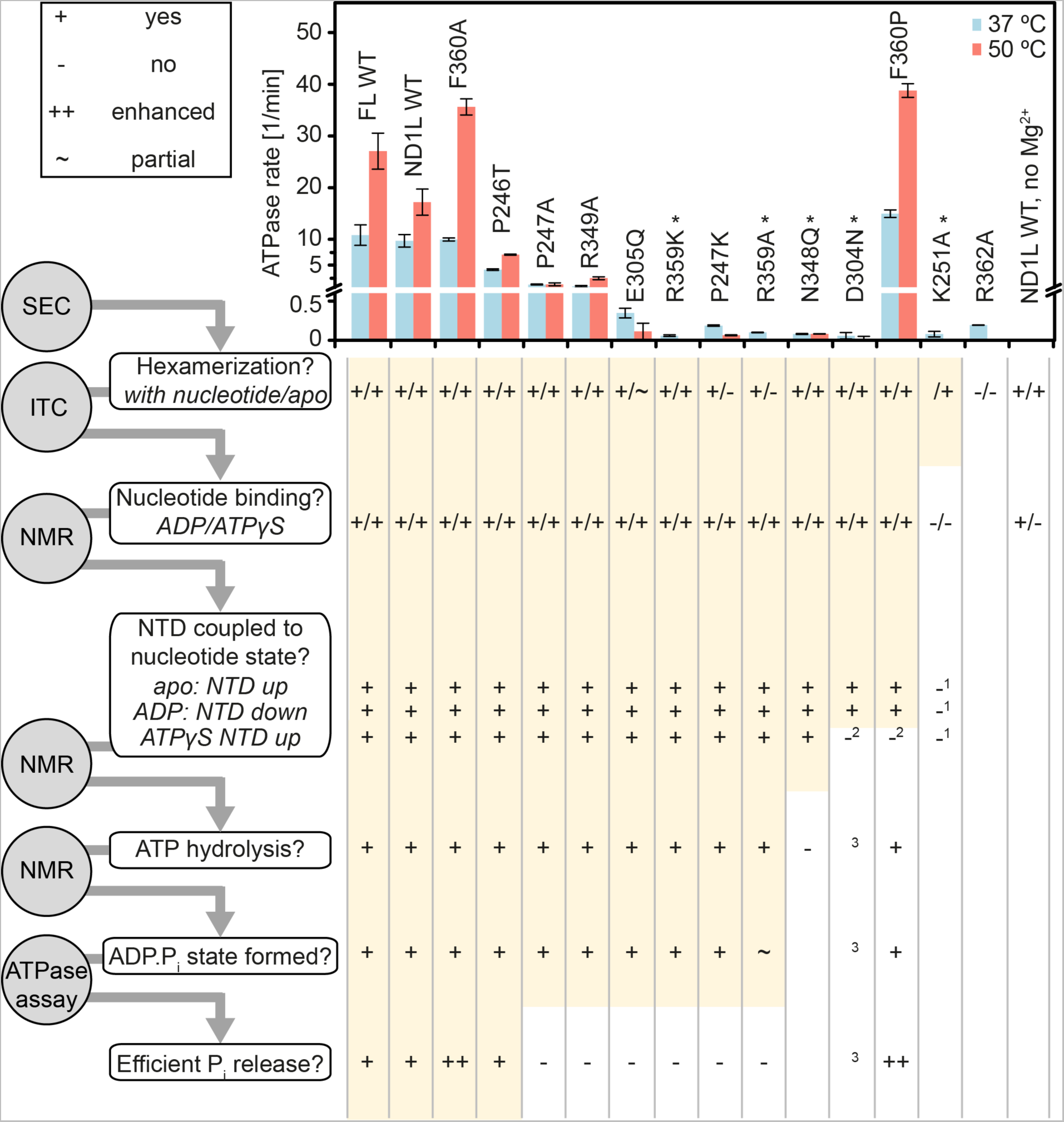
Structural and functional defects of D1-active site mutants. Left: flowchart of biophysical methods to characterize p97-ND1L mutants with respect to their structural and functional integrity. Methods are shown as grey circles: size-exclusion chromatography (SEC), isothermal titration calorimetry (ITC), NMR conformational analysis, ATPase rate measurement; mutant properties are shown as white boxes. Right top: ATPase rates of the mutants. Right bottom: The sequence of assays was pursued as long as a given mutant was assessed positive in the previous category (+, yellow background) but not if it behaved completely (-) or partially (∼) different from the wt. Annotations: * Mutant displays no detectable ATPase activity; ^1^ spectral change detected in the presence of any nucleotide, irrespective of type, NTD position mixed; ^2^ NTD in ‘down’ state in presence of slowly-hydrolysable ATP analogues; ^3^ ADP.Pi state cannot be distinguished from the ATP state by NMR, precluding categorization of the mutant. The corresponding NMR spectra are shown in Fig. E5 and S5-8. The nucleotide dissociation constants are listed in Table S7.

**Extended Data Fig. 5.**
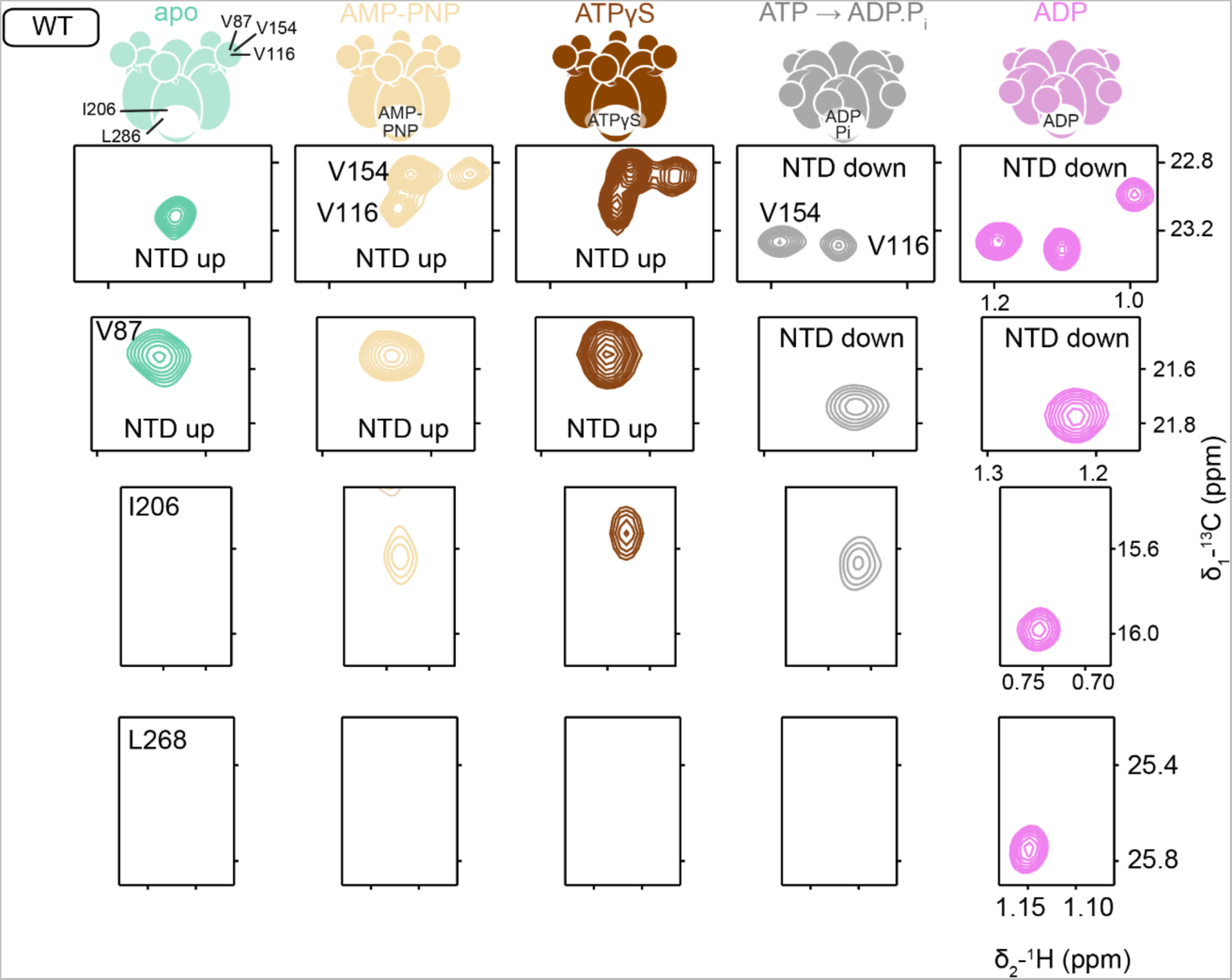
NMR probes indicate global conformation and nucleotide state of p97. Selected spectral regions from HMQC spectra of *proR*-^13^CH3-ILVM-labelled p97-ND1L wt acquired at 37 °C (apo) or 50 °C (all others). Residues V116, V154 and V87 report on the NTD position^15^ (‘up’ in apo, AMP-PNP and ATPγS states *vs*. ‘down’ in ADP and ADP.Pi states), while residues I206 and L268 report on the conformation of the D1 active site and its bound nucleotide. ATPγS^32^ and AMP-PNP^33^ are slowly hydrolysable analogues of ATP. When a mutant assumes the NTD ‘down’ position in the presence of ATPγS, this can be either due to hydrolysis of ATPγS and slow release of thiophosphate or due to a structural defect that prevents the formation of an NTD ‘up’ state in response to Mg^2+^ and ATPγS binding. Therefore, a spectrum in the presence of Mg^2+^ and AMP-PNP, which features a chemically more stable phosphate linkage, was recorded in addition. Excerpts from the corresponding spectra of point mutants (F360P, D304N, K251A, N348Q) are shown in Fig. S5-8.

**Extended Data Fig. 6.**
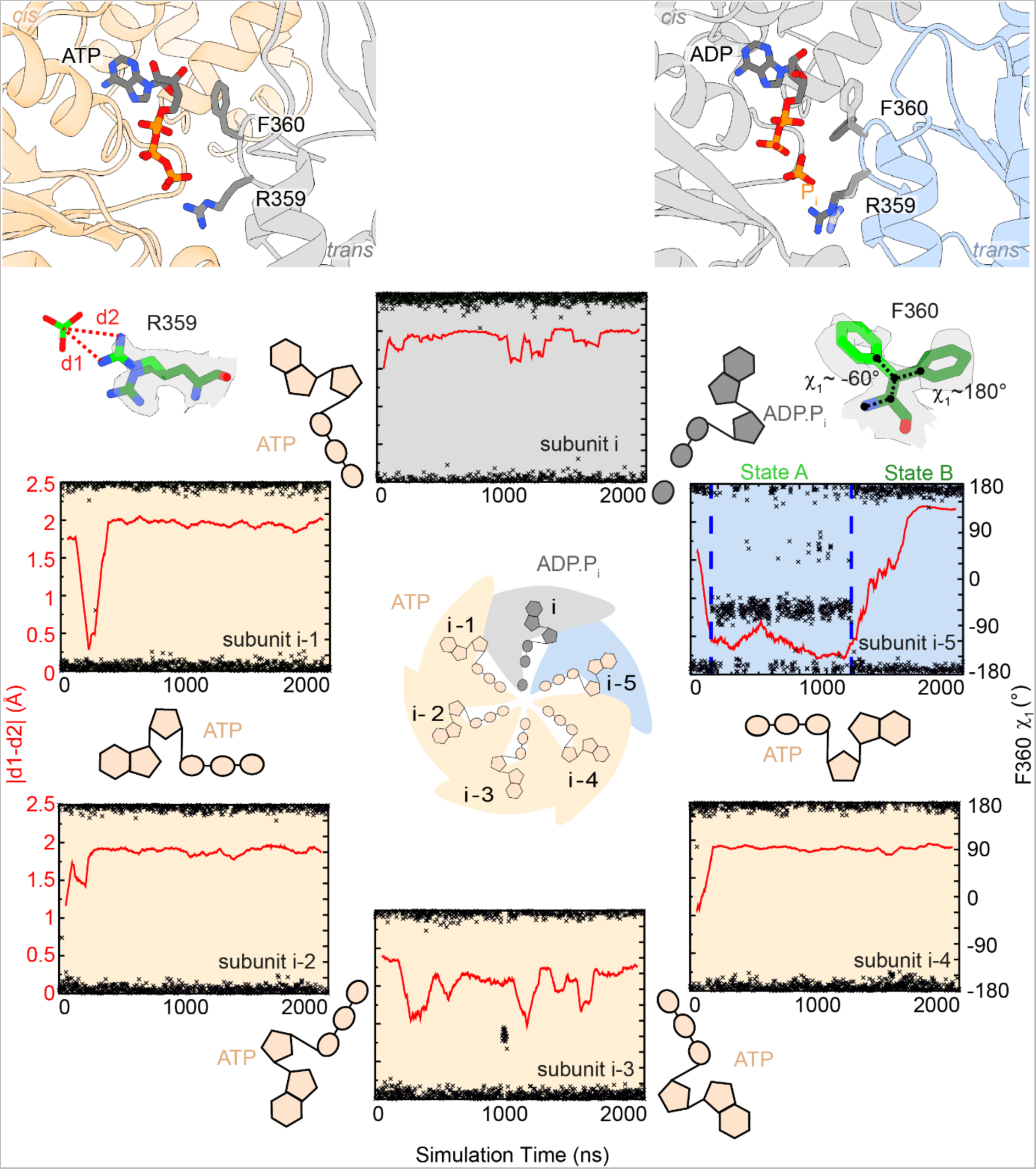
Comparison of active sites in ATP *vs*. ADP.Pi states in MD simulations. The structural and dynamical features of arginine finger residues R359 and F360 at the D1 active site were evaluated over the 2 µs MD trajectory of a p97 hexamer with five active sites occupied with ATP and one with ATP cleaved *in silico* into ADP and Pi. Left ordinate: the distances d1 and d2 between R359-N^ρι^^1^/N^ρι^^2^ and the P atom of the Pi ion reflect the side chain conformation of R359; right ordinate: ξ1 angle of F360. While all ATP-bound subunits show a stable topology throughout, the ADP.Pi-bound subunit undergoes a transition when the Pi ion moves between states A and B, coupled to a flip of the F360 side chain. On top, MD snapshots from the respective active sites are shown highlighting R359 and F360.

**Extended Data Fig. 7.**
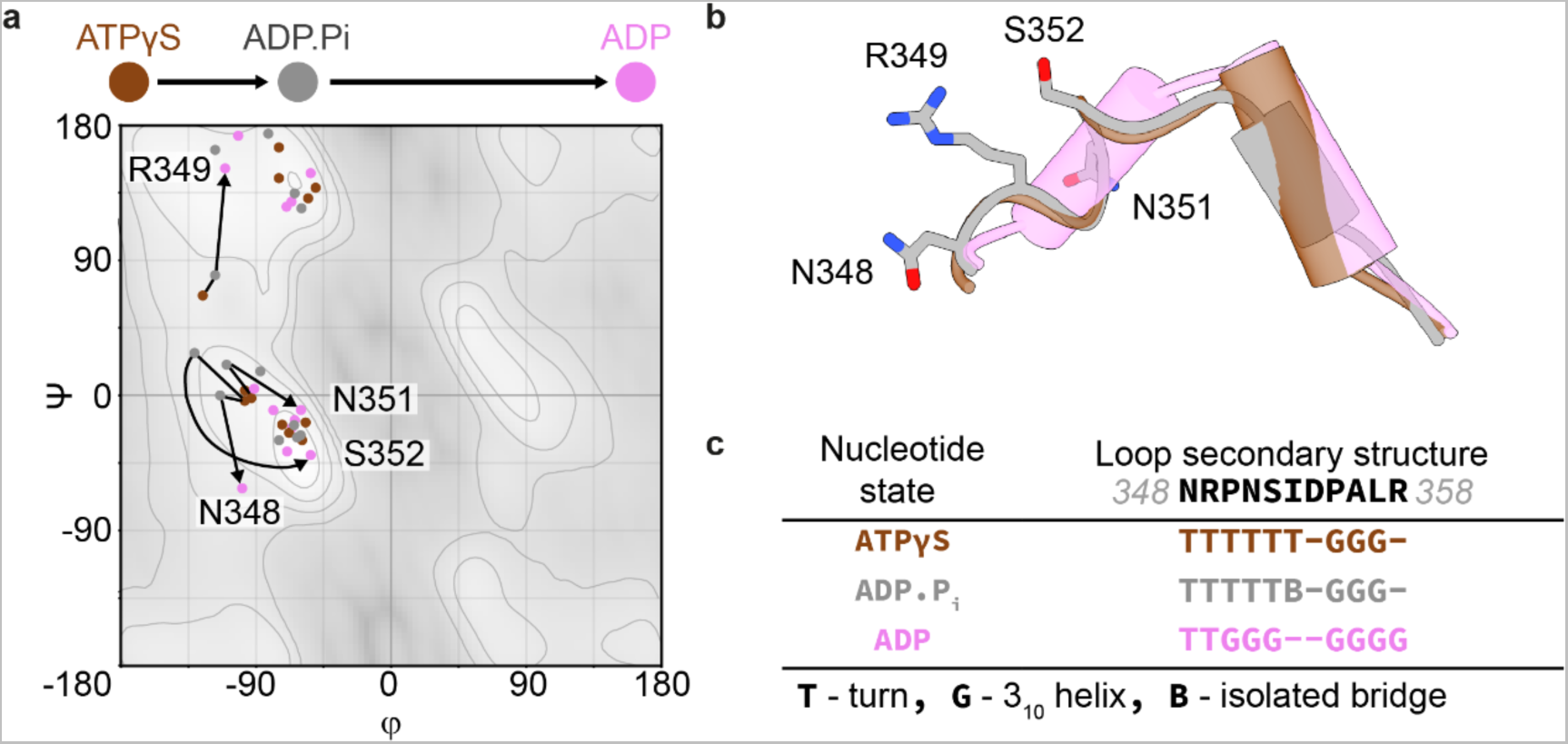
Conformational changes in the sensor loop. **a**, The sensor loop extends from the sensor residue N348 of one subunit to the arginine finger of the adjacent subunit. Its secondary structure is more similar between ATPγS and ADP.Pi states than between ADP.Pi and ADP states, making it the last structural element in D1 to transition from pre-hydrolysis to post-hydrolysis conformation. The transition of the sensor loop is documented by a Ramachandran plot analysis. Residues R359, F360 as well as N348, R349, N351 and S352 undergo dramatic changes in conformation with the progression of ATP-hydrolysis cycle. Residues 349-352 in the first half of the loop change little between ATPγS to ADP.Pi states, but undergo a turn-to-helix conversion from ADP.Pi to ADP state. **b**, Superposition of the sensor loop in different nucleotide states. **c**, For residues 350-352, the STRIDE algorithm^34^ detects a turn for ATPγS and ADP.Pi states and a 310 helix for ADP state. Analyses were performed on the following models: ATPγS: 5ftn^1^, ADP.Pi state A: this work, ADP: 5ftk^1^.

**Extended Data Fig. 8.**
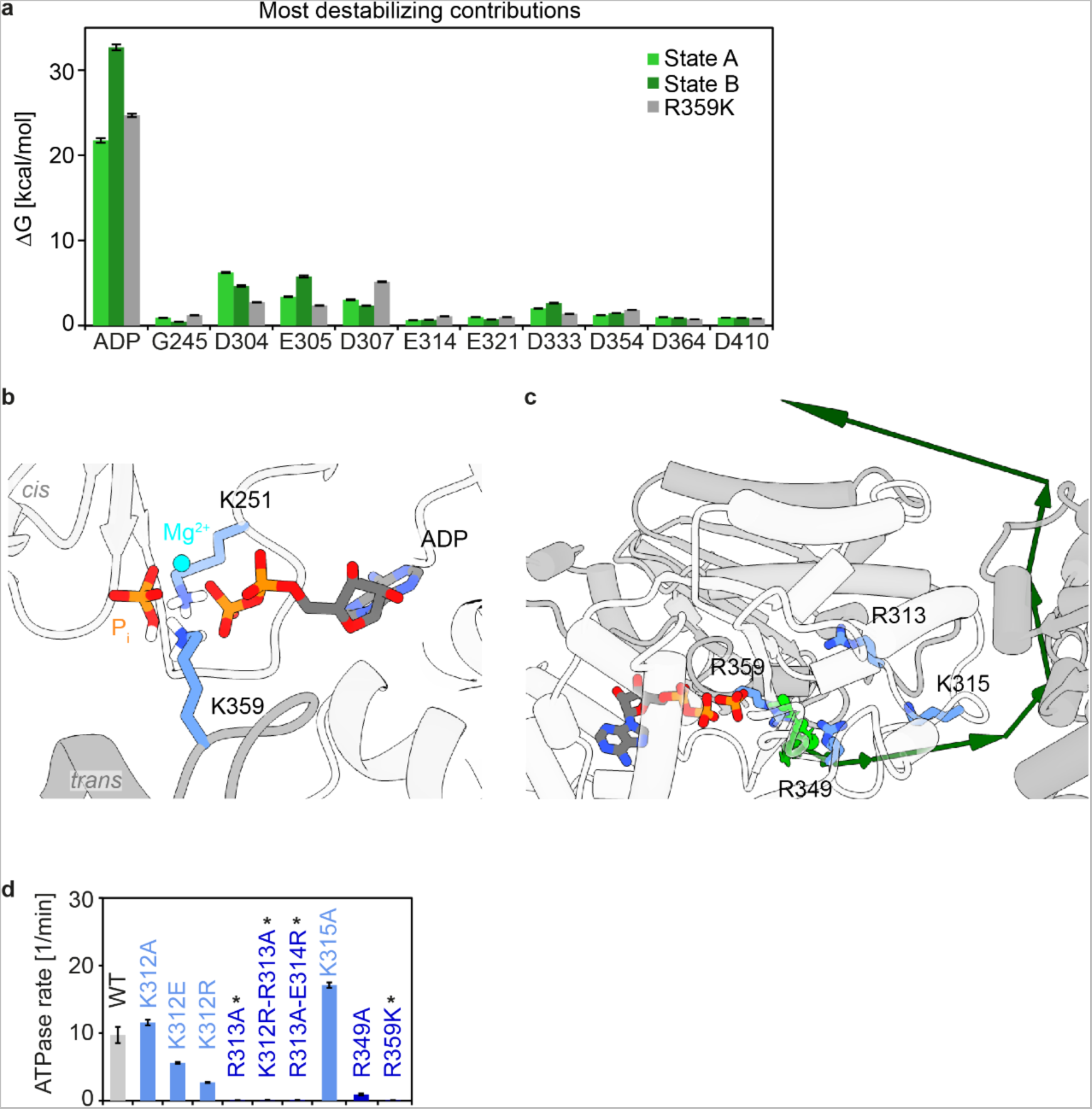
Contributions of active site residues to Pi destabilization and release. **a**, Energy decomposition of MMPBSA calculations identifies residues that destabilize the ADP.Pi states A and B the most. Especially state B is destabilized by the repulsion between ADP and Pi. The overall binding free energy estimated by MMPBSA calculations considers non-bonded interactions (Coulombic, van-der-Waals) between protein and ligand as well as changes in solvation free energy between unbound and bound states. In p97, the most significant factors are polar solvation energies (destabilizing) and electrostatics (stabilizing). Only the electrostatic contributions differ significantly between the calculations for states A, B and R359K mutant. **b**, Lysine side chains contributed by K251 and K359 symmetrically stabilize the ADP.Pi state of the Pi-release deficient mutant R359K. **c**, Side view of the Pi dissociation trajectory (top view in Fig. 5b) shows the Pi ion leaving p97 through the central pore via the top of the hexamer. **d**, ATPase activities of p97-ND1L with mutations of R and K residues that contact the Pi ion during dissociation (Fig. 5d). Note that the removal of R but not K strongly reduces ATP turnover and that the mutation RÒK does not sustain ATPase activity. The ATPase activity also cannot be recovered by introducing R at neighbouring sites (as in K312R-R313A or R313A-E314R).

**Extended Data Fig. 9.**
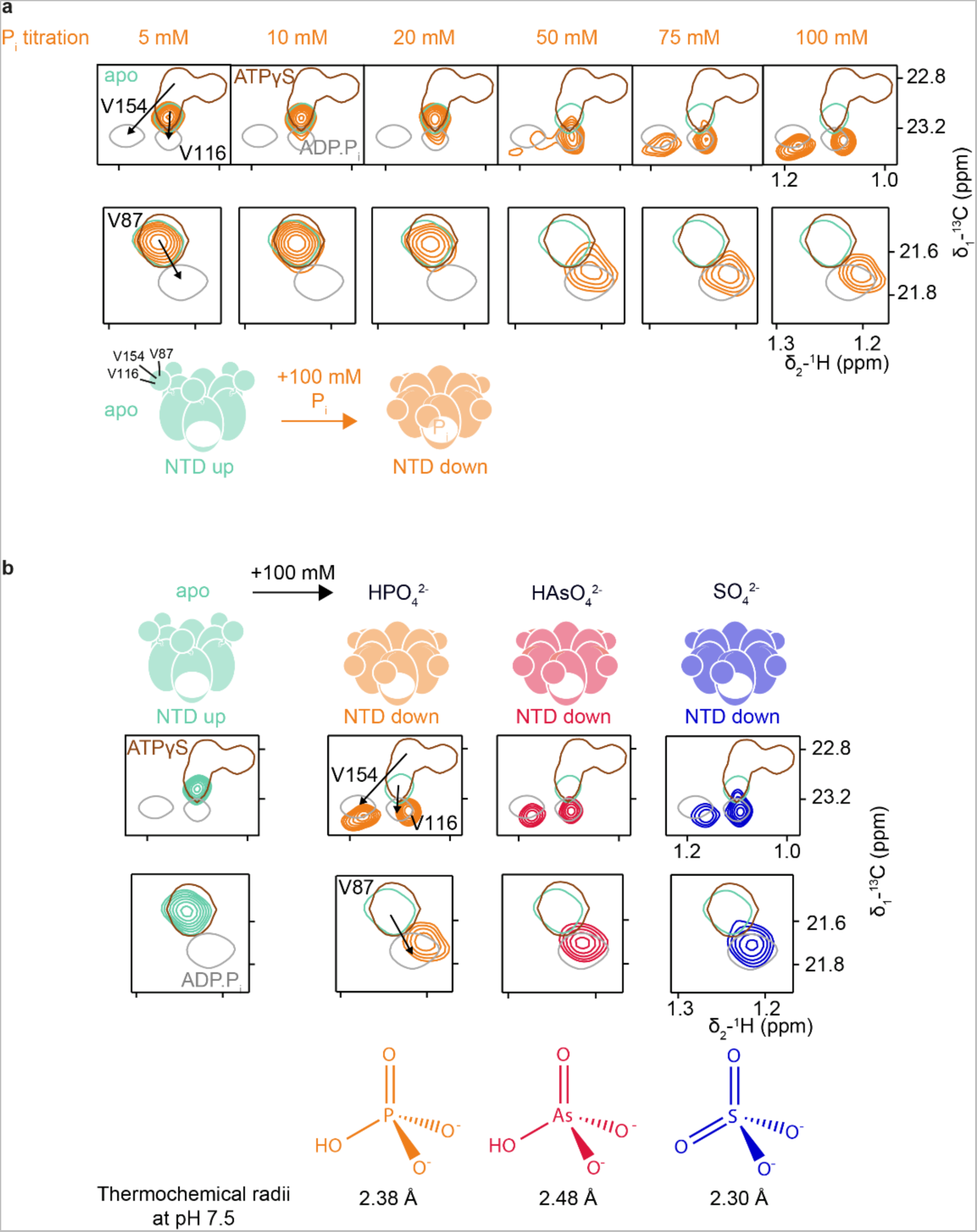
Structural effect of Pi ions and mimics on apo p97. **a**, A concentration of ∼75 mM inorganic Pi ions in solution induces a complete movement of the NTD in apo p97-ND1L into the ‘down’ position. Residues V116, V154 and V87 report on the NTD position. The arrows indicate the shifts of representative methyl correlations from the ‘up’ position (ATPyS state as reference in single contours) to ‘down’ position (ADP.Pi state as reference). Mutants P247K, R359A and R362A do not form hexamers in apo state and do not respond to Pi addition (Fig. S11). Note that fl p97 requires a threefold higher Pi concentration to achieve even a partial effect on NTD position. **b**, Top: Arsenate (HAsO4^2-^) and sulphate (SO4^2-^) can mimic phosphate (HPO4^2-^) at a concentration of ∼100 mM. Bottom: Comparison of thermochemical radii of these ions as an estimate of their effective size in solution.^35^ In the light of our observations, a crystal structure of apo state p97 where the NTD is found in the ’down’ state^36^ may be attributed to a sulphate ion trapped between P247 and K251.

**Extended Data Fig. 10.**
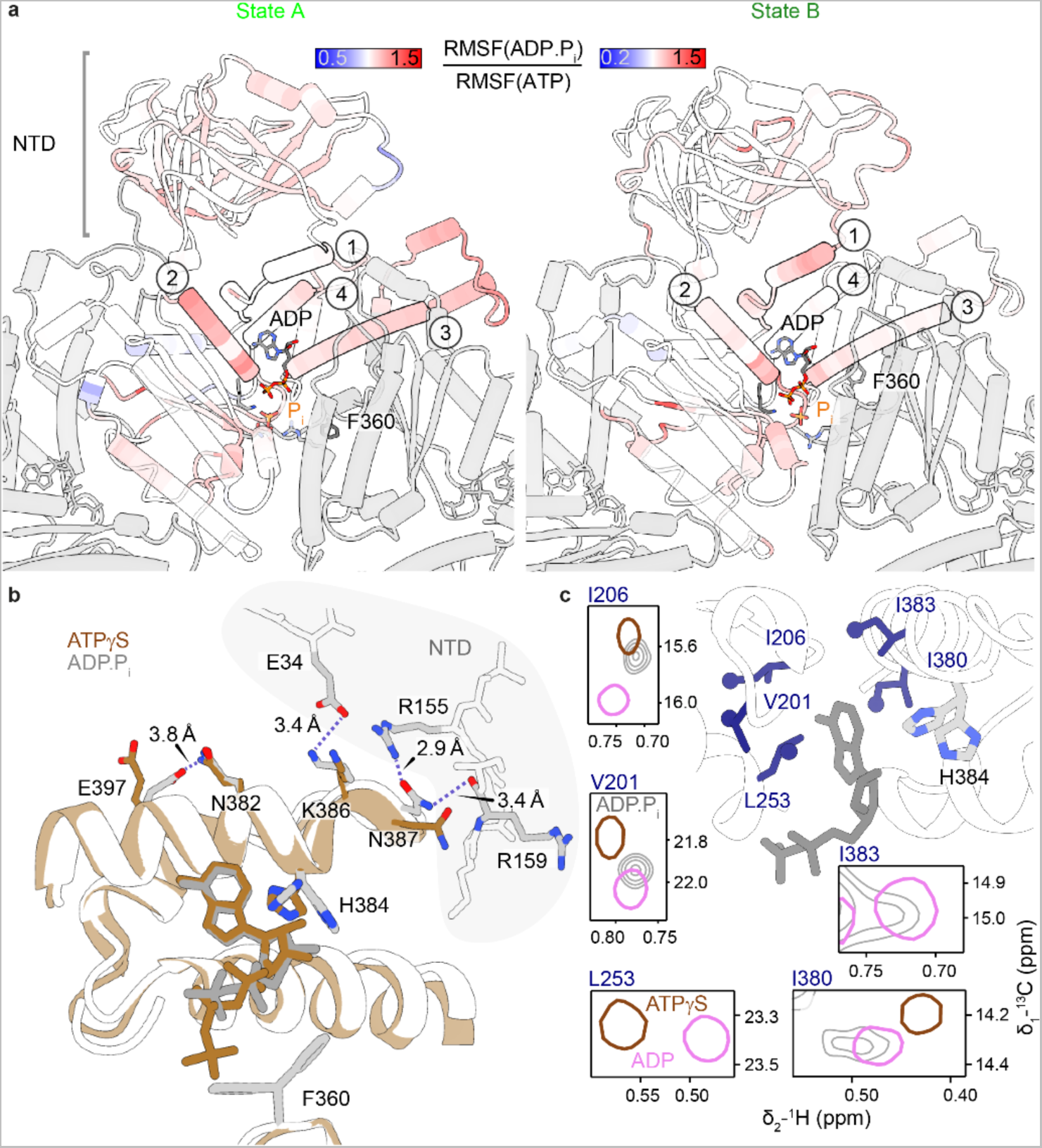
Effects of ATP hydrolysis on protein mobility and structure. **a**, The RMSF of Cα atoms over the 2 µs MD trajectory (same as Fig. E6) quantifies protein mobility at the level of individual residues. The ratio of the RMSF values of the ADP.Pi-bound subunits in states A (left) and B (right) over the average of the five ATP-bound subunits is shown as a heat map on the respective MD snapshots. Increased mobility upon hydrolysis (red shades) is observed propagating from the active site. State B is overall less mobile than state A. Regions of interest include: (1) helix ⍺191-199 and the NTD-D1 linker; (2) helix ⍺251-262 extending from the Walker A motif to the NTD-D1 interface; (3) helix ⍺407-423 to which F360 associates transiently in *trans*; (4) helix ⍺374-387 running past the nucleotide towards the NTD-D1 interface displays increased mobility in state A only. **b**, Two side chain rotamers of H384 are uniquely observed for the ADP.Pi state. They belong to a larger interaction network between D1 and NTD, which enables the conformational change that locks the NTD into the ‘down’ position. In comparison to the ATPγS structure, the ADP position turns 7° in the ADP.Pi structure, enabling an interaction between the ribose moiety and H384. The C-terminus of helix ⍺374-387 forms an interface with the NTD (stick representation) only in the ‘down’ state, where N387 forms a tight network with residues R155 and R159. Meanwhile, N387 flips away from this interface when the NTD is in the ’up’ position (*c.f.* Video S5). **c**, The NMR signals of nearby residues are sensitive both to ATP hydrolysis (ATP𝛾S *vs*. ADP.Pi) and to Pi release (ADP.Pi *vs*. ADP). The location of the methyl probes is visualized on the ADP.Pi state structure.

## Online content

10 extended data figures; supplementary information with the complete experimental methods and references, 12 supplementary figures and 7 tables; 5 supplementary videos.

## Acknowledgement

We thank Riddhiman Sarkar, Pavlo Bielytskyi, Matthias Brandl, Gerd Gemmecker and Sam Asami for support with NMR experiments. This work was funded by the German Research Foundation (DFG) through the Emmy Noether program (project number 394455587 to AKS), SFB1035 (project number 201302640; project B15 to AKS; project A01 to ES; project B02 to MZ), CRC889 (project number 154113120; project A11 to ES), Germany’s Excellence Strategy (project number EXC 2067/1-390729940 to ES). Access to NMR spectrometers was provided by Bavarian NMR Centre of Technical University of Munich and Helmholtz Centre Munich.

## Author contribution

MS, MH and TCC contributed equally. MS, SRR, and YS produced the protein samples. MS performed and analysed NMR experiments and biophysical assays. MH designed, conducted and analysed MD experiments. TCC and SRR performed and analysed cryo-EM experiments. KDL performed and analysed ITC experiments. YS helped devise the protein purification protocol. MZ, ES and AKS designed and supervised the research, MS prepared the figures, AKS and ES wrote the manuscript with contributions from all authors.

## Data availability

Data supporting the findings of this work are available within the Article, Extended Data and the Supplementary Information files. Further details and raw data from *in-silico* modelling are also available from the corresponding authors upon request. Cryo-EM maps, model coordinates and associated structure factors of p97 in ADP.P_i_ states have been deposited in the Electron microscopy Data Bank (EMDB code: 16781/16782) and Protein Data Bank database (PDB code: 8cpf).

## References

1. Banerjee S, Bartesaghi A, Merk A, Rao P, Bulfer SL, Yan Y, et al. 2.3 A resolution cryo-EM structure of human p97 and mechanism of allosteric inhibition. Science 2016, 351(6275): 871–875.

2. Tang WK, Li D, Li CC, Esser L, Dai R, Guo L, et al. A novel ATP-dependent conformation in p97 N-D1 fragment revealed by crystal structures of disease-related mutants. EMBO J 2010, 29(13): 2217–2229.

3. Puchades C, Sandate CR, Lander GC. The molecular principles governing the activity and functional diversity of AAA+ proteins. Nat Rev Mol Cell Biol 2020, 21(1): 43–58.

4. Rydzek S, Shein M, Bielytskyi P, Schutz AK. Observation of a Transient Reaction Intermediate Illuminates the Mechanochemical Cycle of the AAA-ATPase p97. J Am Chem Soc 2020, 142(34): 14472–14480.

5. Taylor EW, Lymn RW, Moll G. Myosin-product complex and its effect on the steady-state rate of nucleoside triphosphate hydrolysis. Biochemistry 1970, 9(15): 2984–2991.

6. Dong Y, Zhang S, Wu Z, Li X, Wang WL, Zhu Y, et al. Cryo-EM structures and dynamics of substrate-engaged human 26S proteasome. Nature 2019, 565(7737): 49–55.

7. Wald J, Fahrenkamp D, Goessweiner-Mohr N, Lugmayr W, Ciccarelli L, Vesper O, et al. Mechanism of AAA+ ATPase-mediated RuvAB-Holliday junction branch migration. Nature 2022, 609(7927): 630–639.

8. Eisele MR, Reed RG, Rudack T, Schweitzer A, Beck F, Nagy I, et al. Expanded Coverage of the 26S Proteasome Conformational Landscape Reveals Mechanisms of Peptidase Gating. Cell Rep 2018, 24(5): 1301–1315 e1305.

9. Flaherty KM, Wilbanks SM, DeLuca-Flaherty C, McKay DB. Structural basis of the 70-kilodalton heat shock cognate protein ATP hydrolytic activity. II. Structure of the active site with ADP or ATP bound to wild type and mutant ATPase fragment. J Biol Chem 1994, 269(17): 12899–12907.

10. Reynolds MJ, Hachicho C, Carl AG, Gong R, Alushin GM. Bending forces and nucleotide state jointly regulate F-actin structure. Nature 2022, 611(7935): 380–386.

11. Heissler SM, Arora AS, Billington N, Sellers JR, Chinthalapudi K. Cryo-EM structure of the autoinhibited state of myosin-2. Sci Adv 2021, 7(52): eabk3273.

12. Sobti M, Ueno H, Noji H, Stewart AG. The six steps of the complete F1-ATPase rotary catalytic cycle. Nat Commun 2021, 12(1): 4690.

13. Caffrey B, Zhu X, Berezuk A, Tuttle K, Chittori S, Subramaniam S. AAA+ ATPase p97/VCP mutants and inhibitor binding disrupt inter-domain coupling and subsequent allosteric activation. J Biol Chem 2021, 297(4): 101187.

14. Tang WK, Xia D. Altered intersubunit communication is the molecular basis for functional defects of pathogenic p97 mutants. J Biol Chem 2013, 288(51): 36624–36635.

15. Schuetz AK, Kay LE. A Dynamic molecular basis for malfunction in disease mutants of p97/VCP. Elife 2016, 5.

16. Priess M, Goddeke H, Groenhof G, Schafer LV. Molecular Mechanism of ATP Hydrolysis in an ABC Transporter. ACS Cent Sci 2018, 4(10): 1334–1343.

17. Wendler P, Ciniawsky S, Kock M, Kube S. Structure and function of the AAA+ nucleotide binding pocket. Biochim Biophys Acta 2012, 1823(1): 2–14.

18. Khan YA, White KI, Brunger AT. The AAA+ superfamily: a review of the structural and mechanistic principles of these molecular machines. Crit Rev Biochem Mol 2021: 1–32.

19. Cooney I, Han H, Stewart MG, Carson RH, Hansen DT, Iwasa JH, et al. Structure of the Cdc48 segregase in the act of unfolding an authentic substrate. Science 2019, 365(6452): 502–505.

20. Miller BR, 3rd, McGee TD, Jr., Swails JM, Homeyer N, Gohlke H, Roitberg AE. MMPBSA.py: An Efficient Program for End-State Free Energy Calculations. J Chem Theory Comput 2012, 8(9): 3314–3321.

21. Jurrus E, Engel D, Star K, Monson K, Brandi J, Felberg LE, et al. Improvements to the APBS biomolecular solvation software suite. Protein Sci 2018, 27(1): 112–128.

22. Erzberger JP, Berger JM. Evolutionary relationships and structural mechanisms of AAA+ proteins. Annu Rev Biophys Biomol Struct 2006, 35: 93–114.

23. Murakami K, Yasunaga T, Noguchi TQ, Gomibuchi Y, Ngo KX, Uyeda TQ, et al. Structural basis for actin assembly, activation of ATP hydrolysis, and delayed phosphate release. Cell 2010, 143(2): 275–287.

24. Merino F, Pospich S, Funk J, Wagner T, Kullmer F, Arndt HD, et al. Structural transitions of F-actin upon ATP hydrolysis at near-atomic resolution revealed by cryo-EM. Nat Struct Mol Biol 2018, 25(6): 528–537.

25. O’Brien MC, McKay DB. Threonine 204 of the chaperone protein Hsc70 influences the structure of the active site, but is not essential for ATP hydrolysis. J Biol Chem 1993, 268(32): 24323–24329.

26. O’Donnell JP, Marsh HM, Sondermann H, Sevier CS. Disrupted Hydrogen-Bond Network and Impaired ATPase Activity in an Hsc70 Cysteine Mutant. Biochemistry 2018, 57(7): 1073–1086.

27. Corbett KD, Berger JM. Structural dissection of ATP turnover in the prototypical GHL ATPase TopoVI. Structure 2005, 13(6): 873–882.

28. Shin M, Puchades C, Asmita A, Puri N, Adjei E, Wiseman RL, et al. Structural basis for distinct operational modes and protease activation in AAA+ protease Lon. Sci Adv 2020, 6(21): eaba8404.

29. de la Pena AH, Goodall EA, Gates SN, Lander GC, Martin A. Substrate-engaged 26S proteasome structures reveal mechanisms for ATP-hydrolysis-driven translocation. Science 2018, 362(6418).

30. Ma W, Schulten K. Mechanism of substrate translocation by a ring-shaped ATPase motor at millisecond resolution. J Am Chem Soc 2015, 137(8): 3031–3040.

31. Twomey EC, Ji Z, Wales TE, Bodnar NO, Ficarro SB, Marto JA, et al. Substrate processing by the Cdc48 ATPase complex is initiated by ubiquitin unfolding. Science 2019, 365(6452).

32. Goody RS, Eckstein F. Thiophosphate analogs of nucleoside di- and triphosphates. Journal of the American Chemical Society 2002, 93(23): 6252–6257.

33. Yount RG, Babcock D, Ballantyne W, Ojala D. Adenylyl imidiodiphosphate, an adenosine triphosphate analog containing a P-N-P linkage. Biochemistry 1971, 10(13): 2484–2489.

34. Heinig M, Frishman D. STRIDE: a web server for secondary structure assignment from known atomic coordinates of proteins. Nucleic Acids Res 2004, 32(Web Server issue): W500–502.

35. Silva JJRFd. The biological chemistry of the elements : the inorganic chemistry of life. Clarendon Press: Oxford, 1991.

36. Hanzelmann P, Schindelin H. Structural Basis of ATP Hydrolysis and Intersubunit Signaling in the AAA+ ATPase p97. Structure 2016, 24(1): 127–139.

