## Supplementary_information for "Unravelling ATP processing by the AAA+ protein p97 at the atomic level"

**Affiliations**

### Supplementary Methods

#### Production of recombinant p97 protein

For NMR experiments, full-length (fl) human p97 (Uniprot P55072) and p97-ND1L (residues 1-480) were produced with N-terminal $His_6$-tag and TEV cleavage site as previously described[1, 2]. Point mutations were introduced using site-directed mutagenesis (New England Biolabs). The following mutants were generated for p97-ND1L: P246T, P247A, P247K, K251A, D304N, E305Q, K312A, K312E, K312R, K312R-R313A, R313A, R313A-E314R, K315A, N348Q, R349A, R359A, R359A-R362A, R359K, F360A, ΔCys-F360C-A413C (ΔCys: C69V-C77V-C105A-C174A-C184W-C209V-C415A), F360P, and R362A; for fl p97: E305Q-E578Q mutant for ssNMR experiments. For cryo-EM experiments, GST-tagged p97 was cloned into a pGEX6p1 vector.

All proteins were over-expressed in *Escherichia coli* BL21(DE3) cells. For solution-state NMR, perdeutration and selective labelling with I-$\delta_1$-[$^{13}CH_3$], V/L-$\gamma_1$/$\delta_1$(*proR*)-[$^{13}CH_3$, $^{12}CD_3$] and M-$\varepsilon_1$-[$^{13}CH_3$] were achieved as previously described[1]. Cells were induced with 0.5-1 mM IPTG 1 hr after the addition of the selective labels and grown over night at 16-18 °C.

$His_6$-tagged p97 constructs were purified[2] using $Ni^{2+}$-NTA affinity chromatography followed by TEV protease cleavage, followed by size exclusion chromatography on Superdex 200 column (Cytiva). Bound nucleotide was removed via apyrase digestion (New England Biolabs) in the presence of 2 mM DTT and 4 mM $CaCl_2$ over night at room temperature, followed by another run on a Superdex 200 column. Protein concentrations were determined photometrically.

GST-tagged fl p97 was bound to GST Sepharose beads (Cytiva). After washing (PBS pH 7.4, 1 mM DTT), p97 was eluted (50 mM Tris pH 8.0, 10 mM glutathione) and subjected to GST tag cleavage by HRV3C protease[3]. The protein was then applied to a Resource Q column (Cytiva) and eluted with a NaCl gradient (50 mM Tris pH 8.0, 0-1 M NaCl), followed by further purification using Superose 6 Increase column (Cytiva) in 25 mM Tris pH 8.0, 150 mM NaCl, 1 mM $MgCl_2$, 0.5 mM Tris(2-carboxyethyl)phosphine (TCEP). Finally, the sample was buffer exchanged to storage buffer (50 mM HEPES, pH 8.0, 150 mM NaCl, 1 mM $MgCl_2$, 0.5 mM TCEP) before snap freezing.

#### NMR sample preparation

For solution-state NMR experiments samples of perdeuterated p97 labelled with *proR*-$^{13}CH_3$-ILVM were buffer exchanged (25 mM HEPES pH 7.5, 25 mM NaCl, 5 mM TCEP, 100% $D_2O$) to concentrations in the range of 50-200 μM. For assessment of the different nucleotide states, the protein samples were supplemented with 5 mM ADP or 4 mM $MgCl_2$ and 5 mM ATPγS or AMP-PNP (Jena Bioscience). Setup of the ATP regeneration system was achieved as previously described[1]. For solid-state NMR measurements, 3 mg of ND1L-E305Q or fl-E305Q-E578Q at natural isotopic abundance were dialyzed (25 mM HEPES pH 7.0, 50 mM NaCl, 5 mM TCEP, 100% $H_2O$), supplied with the regeneration system and sedimented into 1.3 mm MAS rotors (Bruker) using filling tools (Giotto Biotech).

#### NMR titrations

The apo state of p97-ND1L wt was titrated with inorganic phosphate ($P_i$) in several steps from 0 up to a final concentration of 100 mM from an 800 mM $Na_2HPO_4$ pH 7.5 stock solution. All mutants were supplied with inorganic $P_i$ to a concentration of 100 mM in one step.

Mimics of $P_i$ ions were added to the apo state of p97-ND1L wt in one step to a final concentration of 100 mM (stocks: $Na_2HAsO_4$ dissolved to 300 mM; $Na_2SO_4$ dissolved to 1 M; stocks adjusted to pH 7.5).

#### NMR spectroscopy

Solution-state NMR experiments were conducted on Avance III Bruker spectrometers equipped with TCI cryo probes at field strengths corresponding to proton resonance frequencies of 800, 900 and 950 MHz. Sample temperatures during data acquisition were 37 °C (all apo states), 40 °C (all K251A spectra; F360A/P spectra in the presence of ATP) or 50 °C (all others).

Solid-state NMR experiments were performed on an Avance III 800 MHz Bruker spectrometer under 45 kHz MAS at 5 °C[1]. Cross-polarisation based experiments were measured interleaved with directly pulsed experiments for reaction control. Chemical shifts were referenced to internal sodium trimethylsilylpropanesulfonate (DSS, Sigma).

Experimental parameters are listed in Tables S1 and S2. All spectra were processed using TopSpin (Bruker; V 3.5) and analysed using CcpNmr Analysis (CCPN, V 2.4.2)[4]. The $^{31}P$ spectrum (Fig. 1b) was fitted using Mnova 11.0 (Mestrelab).

### Cryogenic electron microscopy

#### Grid plunging and cryo-EM data acquisition

Purified p97 was concentrated to ~4 mg/ml and incubated in the ATP regeneration system (4 mM ribose-5-phosphate, 4 mM $MgCl_2$, 50 mM KCl, 13.3 U pyruvate kinase, 50 mM phospho-enol pyruvate, 10 mM ATP) for 20 mins at 37 °C. Octyl-beta-glucoside at a concentration of 0.05 % was added just before plunge freezing. 3 μl of sample was blotted on glow discharged Quantifoil Cu R2/1, 200 mesh grids. Plunge freezing was performed with a Vitrobot Mark IV (Thermo Fisher Scientific) in a chamber equilibrated at 10 °C with 100 % humidity. Images were acquired with a Titan Krios G4 (Thermo Fisher Scientific), with a Falcon 4 detector (Thermo Fisher Scientific) mounted after a Selectris energy filter with slit width at 15 eV. A total of 10011 images were collected using EPU with aberration free image shift (AFIS), at 165kx magnification (0.72 Å/pix). Each image had an exposure of 40 e/$Å^2$, with an exposure rate of 5.41 e/pix/s. The nominal defocus range was from -0.9 to -2.2 μm.

#### Cryo-EM image processing

Each image consisted of 931 electron-event representation (EER) frames. Initial drift correction was performed with Motioncorr[5] as implemented within Relion[6], such that a grouping of 23 EER frames were used. Contrast transfer function (CTF) parameters were estimated by CTFFIND4[7]. A total of 693,686 particles were picked with Cryolo[8] using a p97-trained network. Relion 4.0[9] was used for subsequent data processing. All particles were first extracted in bin2 and subjected to initial cleaning up by 2D and 3D classification. 199,453 double ring p97 particles were selected and re-extracted without binning in a box of 330 pixels for further processing, and 3D refinement reached a global resolution of 2.98 Å after CTF, magnification and higher optical aberration correction. In order to obtain a high-resolution map for in-depth analysis of the D1-D2 domain, Bayesian polishing was performed and followed by subtraction of single-ring from double-ring particles, which effectively doubled the total number of particles to 398,906 and led to a reconstruction of 2.83 Å with C6 symmetry applied. A subset of 181,651 particles were identified by a focused 3D classification without alignment, which produced a 2.64 Å map (C6 symmetry applied) after CTF and higher optical aberration correction. As a final push of resolution, a limit to use only particles with closer defocus than -1.7 μm reduced the number of particles to 86,760 that yielded a map of 2.61 Å (C6 symmetry applied).

Because the NTD domain is typically more flexible than the D1-D2 ring, the following image processing was performed to improve the map quality of the NTD domain. The 181,651 computationally subtracted single rings were reverted back to double rings, and duplicated particles were removed such that 125,454 particles remained. Subtraction of single-ring from double-ring particles yielded a total of 250,908 particles. After further 3D refinements and 3D classification without alignment, 112,231 particles were selected and a signal subtraction was performed to focus on only one p97 subunit, which finally yielded a map of 3.27 Å with sufficient NTD density quality for interpretation. Acquisition parameters and statistics are listed in Tab. S3.

#### Initial cryo-EM model building

The published models PDB 5ftm and 5ftl[10] were used as a starting point for model building. The model was first rigid-body docked to the D1-D2 ring focused map and NTD focused map, followed by manual adjustment in Coot[11]. The model was then refined by phenix.real_space_refine. A composite whole map of p97 was constructed by combing the two focused refined maps of D1-D2 ring and NTD using phenix.combine_focus_maps[12]. The models were docked into the composite maps to generate a complete model of p97 (see Table S4). The model was then subjected to molecular dynamics simulation for the analysis of $P_i$ and $Mg^{2+}$ ion positions.

#### Molecular dynamics simulations

Molecular dynamics simulations were performed using the GPU accelerated version of pmemd[13], as distributed with the AMBER18[14] package. For proteins the ff14Sb[15] force field was used while water molecules were described with the SPC/E[16] model. In order to model the crucial $Mg^{2+}$ ions as accurately as possible, the 12-6-4LJ model[17] for divalent ions was used. ATP and ADP parameters[18] were taken from the parameter database of the University of Manchester. Single and double protonated $P_i$ ions ($HPO_4^{2-}$ and $H_2PO_4^-$) were parameterized utilizing the GAFF2[19] force field for organic molecules. For this RESP[20] partial charges (Hartree-Fock at 6-31G* level) were calculated from a structure that was optimized to a gas-phase energy minimum at the B3LYP/TZVP-level. All QM calculations were conducted with GAUSSIAN09[21].
All simulations were performed at 303.15 K and a pressure of 1 atm using the Langevin[22] thermostat (with a collision frequency of 1 $ps^{-1}$) and the Monte Carlo barostat[23] ($\tau_p$ = 1.0 ps), respectively.
Non-bonded interactions were calculated explicitly until a distance cut-off of 9 Å. Long-range Coulomb interactions were accounted for by the particle mesh Ewald method[24], while long-range van-der-Waals effects were described by a dispersion correction model. During the sampling phase, time steps of 4.0 fs were used, which was enabled by constraining all bonds involving hydrogen atoms[25] to their equilibrium lengths as well as applying the hydrogen mass repartitioning method[26].
Data analysis was performed using VMD 1.9.3[27] and CPPTRAJ[28].

##### Simulation setup

The molecular dynamics simulations of wild-type (wt) p97-ND1L in ADP.$P_i$ state and point mutations thereof (N348Q, R359K, F360P) were started from the crystal structure with PDB ID 4ko8 [29], which lacks the D2 subunit. Hexamers were generated from the asymmetric unit that contains a dimer encompassing residues 14-469. ATP$\gamma$S was transformed into ATP (in five of six subunits) while ADP and either $H_2PO_4^-$ or $HPO_4^{2-}$ were placed in one subunit. The starting position of the *in-silico* created $P_i$ ion was chosen so that it coincides with the position of the former $\gamma$-phosphate.

Protonation states of titratable groups were assigned using the PDB2PQR server (https://server.poissonboltzmann.org/pdb2pqr)[30] at pH 7.4. Only the protonation state of Lys251 was set by hand (to charged). The simulations were performed using periodic boundary conditions in octahedral simulation boxes containing approx. 117,000 water molecules as well as 25 mM NaCl and 50 mM KCl.

#### Final model building

The simulation conducted for the refinement of the cryo-EM structure was started from a preliminary cryo-EM structure of fl p97. All six D1 binding sites contained ADP + $HPO_4^{2-}$ + $Mg^{2+}$, all six D2 binding sites contained ATP + $Mg^{2+}$. The solvent consisted of approx. 106000 water molecules and $Na^+$ counter ions. Periodic boundary conditions were applied. Unexplained cryo-EM densities around the D1 pocket were explained by the simulation. In the final model, two states, A and B, of dissociating $P_i$ ions with their corresponding $Mg^{2+}$ ions were identified.

The simulation sampling $P_i$ ion dissociation was performed under identical conditions except that the $Mg^{2+}$ ions bridging ADP and $HPO_4^{2-}$ were removed from the system.

An overview over all conducted simulations is given in Table S5.

Prior to sampling a seven-step equilibration protocol was applied to all simulation systems (details in Table S6).

#### Free energy calculations

The interaction free energies (ΔG) of $P_i$ and ND1L-p97 (wt in states A and B as well as the mutant R359K) were calculated using the MMPBSA single-trajectory-method as implemented in the MMPBSA.py[31] script, which is part of the AMBER18 package[14]. Since the Poisson Boltzmann (PB) routine of AMBER18 was unable to recognize one of the atom types in the ATP force field, the PB calculations were performed with flags inp=1 and radiopt=0, which resulted in slightly different nonpolar solvation terms compared to the default settings in AMBER18.

The numbers of processed frames for each system were as follows: wt state A: 450 frames, wt state B: 400 frames, R359K: 500 frames. A salt concentration of 150 mM was chosen. The dielectric constant for the protein was set to 1.0, while water was set to 80.0.

Conformational entropy contributions were neglected because of the high computational cost and generally low accuracy of these methods. Therefore, the resulting ΔG values cannot be directly compared to experimental values. However, our analyses compare very similar systems (slightly different conformations of the same protein and a point mutant thereof), therefore it can be assumed that errors stemming from this treatment cancel each other to a very high degree.

### Biochemical assays

#### Intersubunit crosslinking

To apo state p97-ND1L-ΔCys-F360C-A413C, 4 mM DTT were added and the solution was incubated for 2 h at 30 °C. Reducing agent was removed by gel filtration on a Superdex 200 in crosslinking buffer (20 mM HEPES pH 7.2, 250 mM KCl, 5 mM EDTA). Fractions eluting as hexamers were pooled and diluted to 20-60 μM. Crosslinking reagent bismaleimidoethane (BMOE, ThermoFisher Scientific) was supplied in twofold excess from a 20 mM stock in DMSO and the solution was incubated for 2 h on ice. After quenching with 50 mM DTT (15 min on ice), excess chemicals were removed via another gel filtration run on a Superdex 200 in gel filtration buffer. Only protein eluting as hexamer was pooled for further studies. Successful crosslinking was verified by SDS-PAGE (Fig. S12).

#### Size-exclusion chromatography (SEC)

The oligomerisation states of the various p97 mutants were estimated from the elution profile following SEC on a Superdex 200 Increase 10/300 GL gel filtration column (Cytiva; buffer: 50 mM HEPES pH 7.5, 250 mM KCl, 2 mM $MgCl_2$). Size calibration was achieved internally using molecular weight standards (SERVA).

#### NADH-coupled ATPase assay

The ATPase rates of p97 were determined using an NADH-coupled ATPase assay, where the oxidation of NADH is directly coupled to the rate of ATP hydrolysis. Phosphoenolpyruvate (6 mM), NADH (1 mM), pyruvate kinase (1U/100 μL), lactose dehydrogenase (1U/100 μL) and purified protein (1-100 μM) were diluted into the ATPase buffer (25 mM HEPES pH 7.5, 25 mM NaCl, 50 mM KCl, 4 mM $MgCl_2$, 0.5 mM TCEP) and distributed into a 96 well plate to a final volume of 120 μL. The reaction mixture was equilibrated at either 37 °C or 50 °C for 5 min before the addition of ATP (2 mM). The decrease in absorbance at 340 nm (coupled to NADH oxidation) was monitored with a SpectraMax iD5 plate reader (Molecular devices) for 60 min. Data were analysed using in-house scripts. ATP-hydrolysis rates (ATP/min) were calculated based on at least three experimental replicates.

#### Isothermal titration calorimetry (ITC)

ITC measurements were conducted on a MicroCal PEAQ-ITC (Malvern Pananalytical) instrument at 25 °C. Protein samples (10–20 μM) were freshly digested with apyrase as described above and subsequently run over a Superdex 200 column (Cytiva); lyophilized commercial nucleotides (100-120 μM; Sigma and Jena Bioscience) were dissolved in identical SEC buffer. Experimental parameters included one 0.4 μL injection followed by 19x2 μL injections with 120 s of spacing. Data were analyzed using MicroCal PEAQ-ITC Analysis software (V 1.21) using the One Set of Sites binding model. $K_d$ values were calculated based on at least two experimental replicates.

#### Electrostatic potential calculation

Electrostatic potential calculations were performed using the Adaptive Poisson-Boltzmann Solver (APBS)[30] with input generated from the cryo-EM derived structure (state A) with amendments from the PDB2PQR program[32].

#### Sequence alignment

All multiple sequence alignments were done using Clustal Omega[33]. Sequences of AAA+ ATPases having two tandem ATPase domains such as NSF and p97 were edited in Jalview[34] prior to the domain alignment.

Accession IDs of the sequences used for human AAA+ ATPases alignment (Fig. 3a):

P35998 (PSMC2), P46459 (NSF), Q16740 (CLPP), P36776 (LONP1), Q13608 (PEX6), Q9Y265 (RUVBL1)

Accession IDs of the sequences used for alignment of p97 from different species (Fig. S4):

Q7KN62 (TER94_*D. melanogaster*), Q9P3A7 (CDC48_*S. pombe*), P25694 (CDC48_ *S. cerevisiae*) P46462 (VCP_ *R. norvegicus*), Q3ZBT1 (VCP_ *B. taurus*), P54812 (CDC48.2_*C. elegans*), Q01853 (VCP_*M. musculus*), P55072 (VCP_*H. sapiens*)

#### Ramachandran Plot Analysis

RamachanDraw (https://github.com/alxdrcirilo/RamachanDraw) was used to create Ramachandran plots. Torsion angles of residues 348-360 in the cryo-EM structure of ADP.$P_i$ state (this work), ATPγS state (PDB ID 5ftn[10]), and ADP state (PDB ID 5ftk[10]) were considered in this analysis.

#### Visualization

Molecular graphics and analyses were performed with UCSF Chimera 1.16[35] and ChimeraX 1.4[36].

### Supplementary Figures

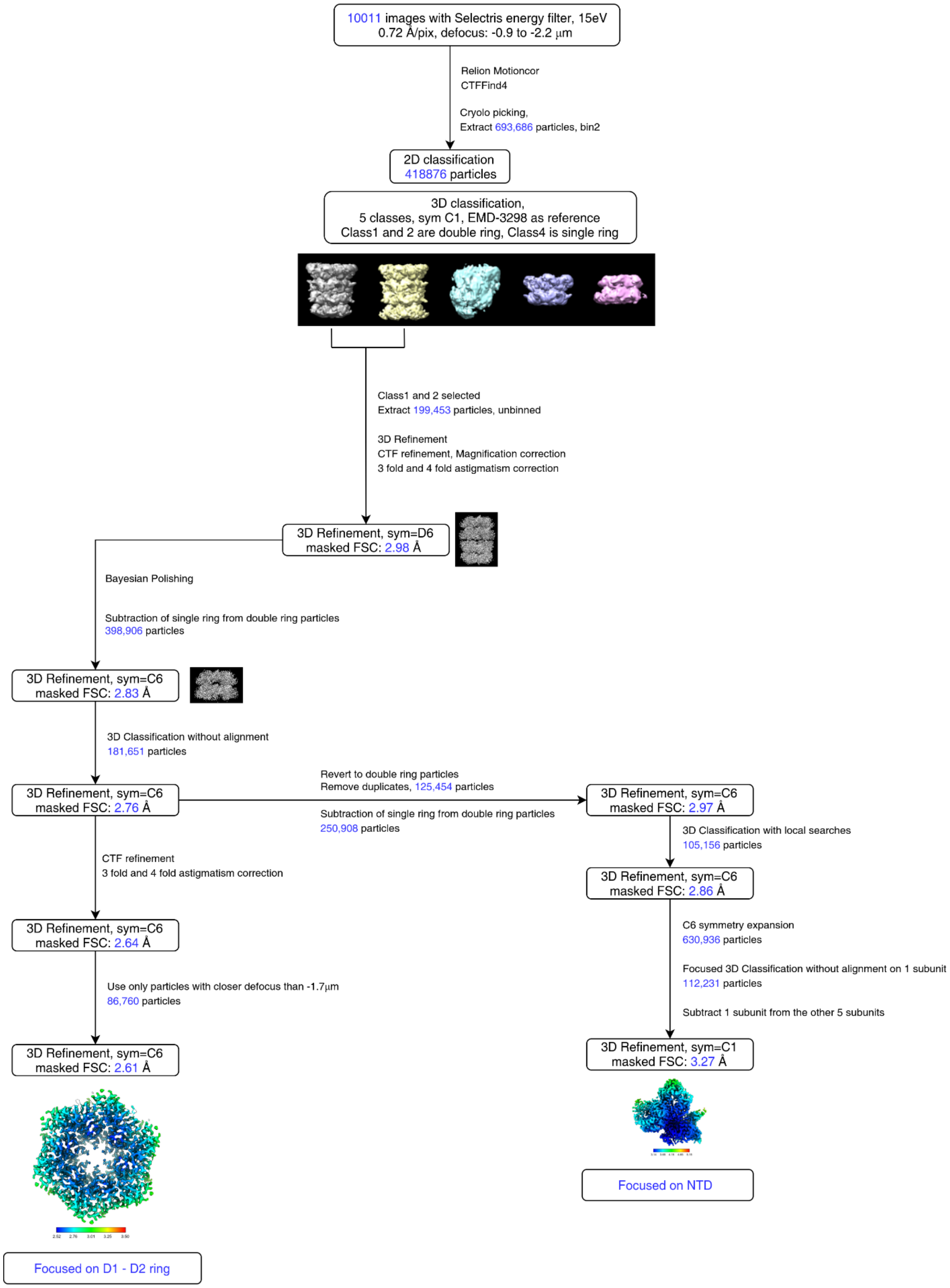

**Supplementary Fig. 1 Flow chart of cryo-EM data analysis.** Reconstruction of wt fl p97 in the presence of an ATP-regeneration system. Classification and refinement procedures for the D1-D2 focused maps and NTD focused maps are shown.

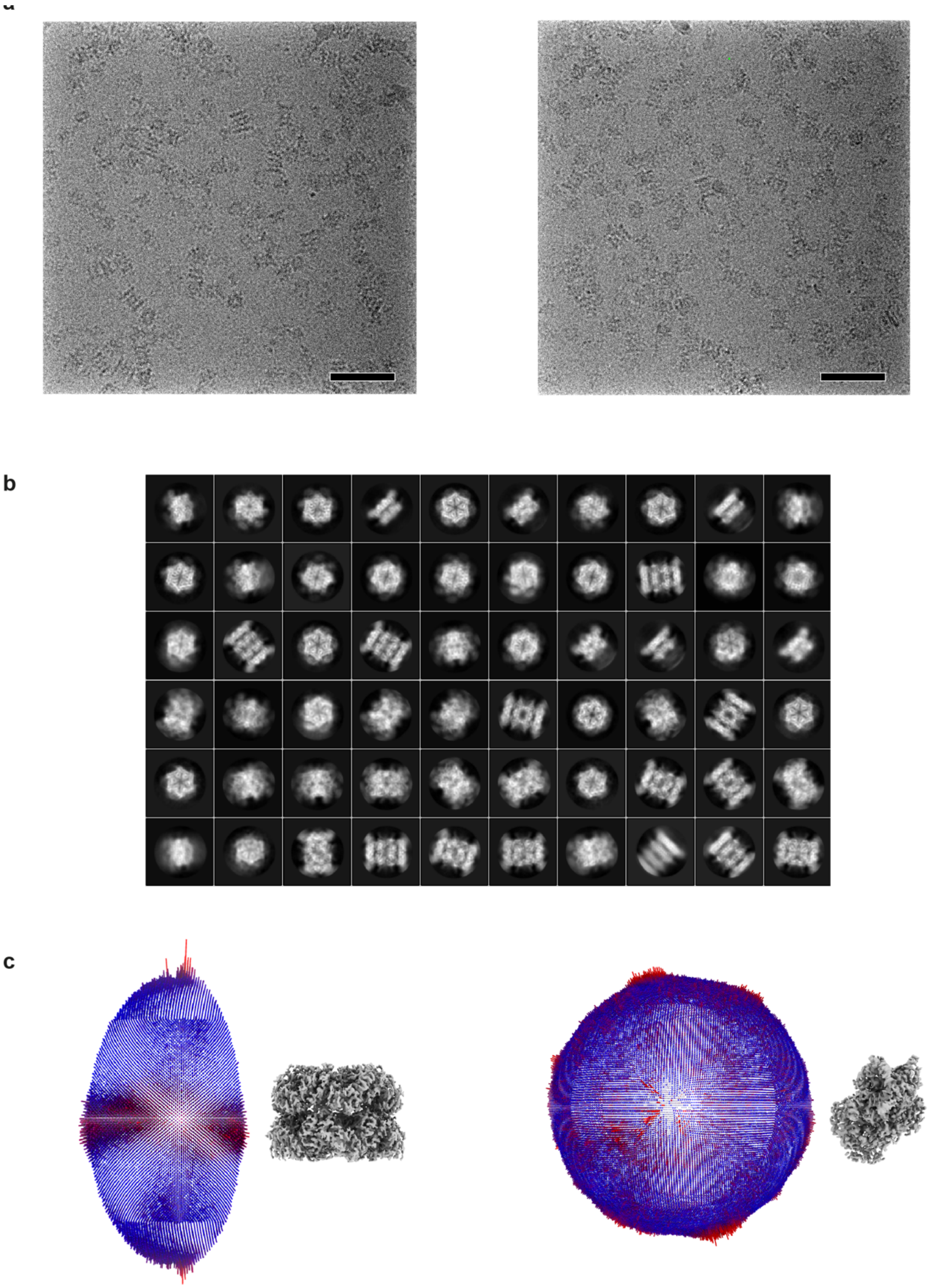

**Supplementary Fig. 2 Cryo-EM image processing of p97 in the ATP regeneration system.** **a**, Representative micrographs of wt p97 in the presence of an ATP-regeneration system, scale bar represents 50 nm. **b**, Two-dimensional class averages. **c**) Angular distribution of particle views for the reconstruction with focus on the D1-D2 ring (left) and with focus on the NTD (right).

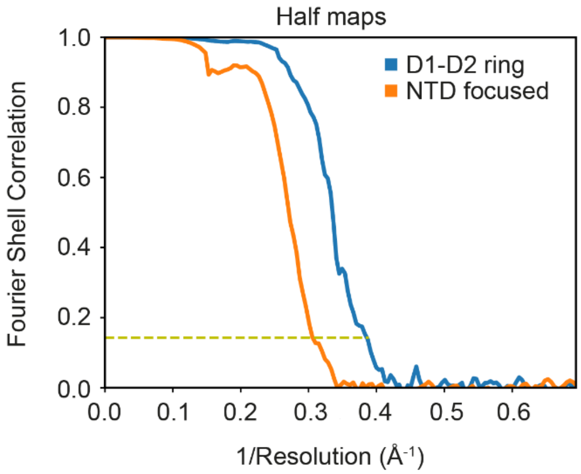

**Supplementary Fig. 3 FSC curves of cryo-EM reconstructions.** Resolutions were determined on the basis of the gold standard Fourier shell correlation between two independently refined half maps[37] (FSC=0.143, dotted line). The D1-D2 ring focused map has a resolution of 2.61 Å compared to 3.27 Å for the NTD focused map.

Seqence alignment in D1 domain

| | | | | | | | | | | | | |
|---|---|---|---|---|---|---|---|---|---|---|---|---|
| *C. elegans* | CDC48 | 250 | GPPGTGKT | 257 | -- | 305 | ILFIDE | 310 | -- | 353 | NRPNSIDGALRRFGR | 367 |
| *D. melanogaster* | TER94 | 242 | GPPGTGKT | 249 | -- | 297 | IIFIDE | 302 | -- | 345 | NRPNSIDPALRRFGR | 359 |
| *B. taurus* | p97 | 245 | GPPGTGKT | 252 | -- | 300 | IIFIDE | 305 | -- | 348 | NRPNSIDPALRRFGR | 362 |
| *H. sapiens* | p97 | 245 | GPPGTGKT | 252 | -- | 300 | IIFIDE | 305 | -- | 348 | NRPNSIDPALRRFGR | 362 |
| *M. musculus* | p97 | 245 | GPPGTGKT | 252 | -- | 300 | IIFIDE | 305 | -- | 348 | NRPNSIDPALRRFGR | 362 |
| *R. norvegicus* | p97 | 245 | GPPGTGKT | 252 | -- | 300 | IIFIDE | 305 | -- | 348 | NRPNSIDPALRRFGR | 362 |
| *S. cerevisiae* | CDC48 | 255 | GPPGTGKT | 262 | -- | 310 | IIFIDE | 315 | -- | 358 | NRPNSIDPALRRFGR | 372 |
| *S. pombe* | CDC48 | 265 | GPPGTGKT | 272 | -- | 320 | IIFIDE | 325 | -- | 368 | NRPNSIDPALRRFGR | 382 |

**Supplementary Fig. 4 Sequence alignment of p97 in different eukaryotic organisms.** The phenylalanine residue between the arginine fingers in D1 (F360 in human p97) is a special feature not found in other eukaryotic AAA+ proteins (cf. Fig. 3a). While it is strictly conserved among p97 homologues, this position is often occupied by a proline residue in AAA+ proteins.

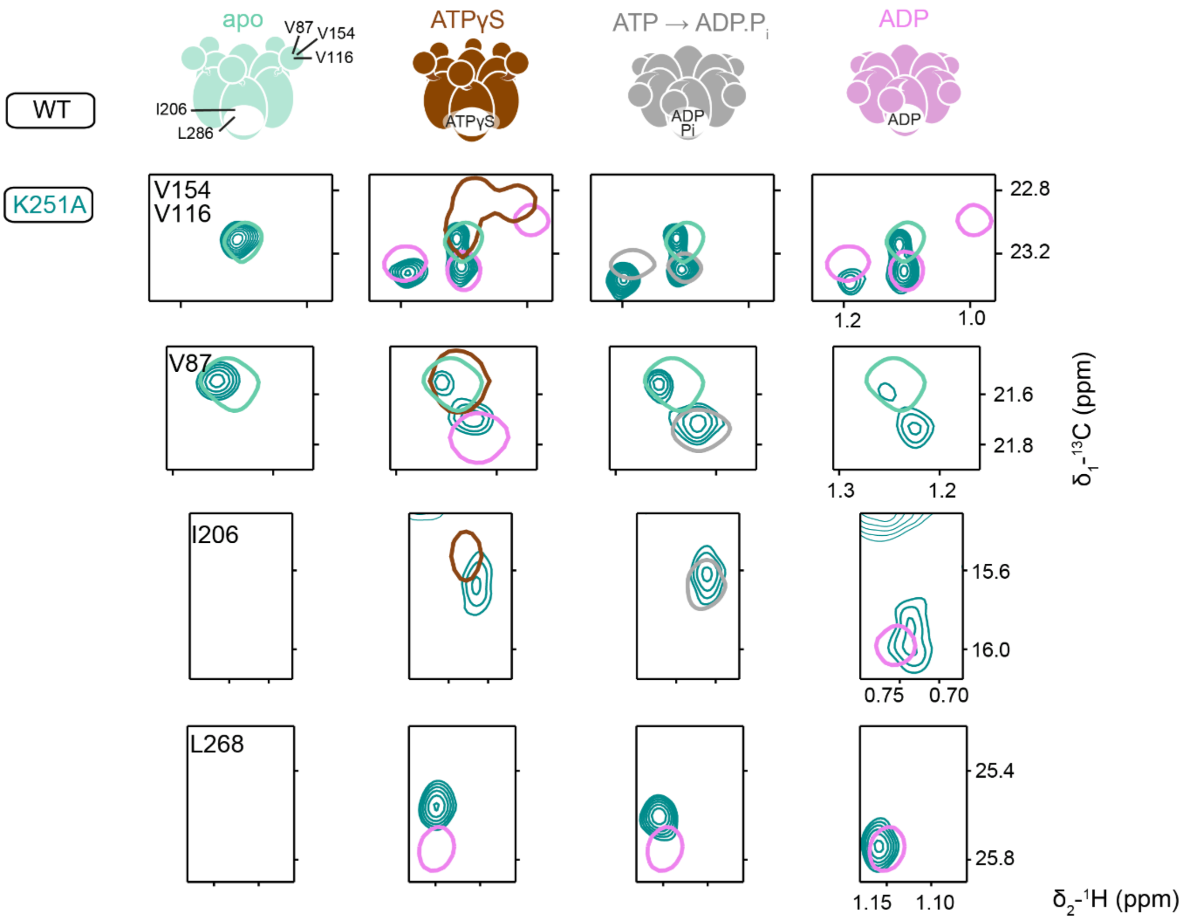

**Supplementary Fig. 5 NMR spectra of p97-ND1L K251A.** Selected spectral regions from ($^{1}H$,$^{13}C$)-HMQC spectra of p97-ND1L K251A as a function of nucleotide present. Residues V116, V154 and V87 report on the NTD position, while residues I206 and L268 reflect the conformation of the D1 binding pocket. Methyl group correlations from the respective wt spectra are indicated in single contours; when wt and mutant spectra differ, the corresponding wt-ADP spectra are shown in addition. Although the mutant lacks the K251 residue that is critical for nucleotide binding[38], the NMR spectra are still sensitive to the presence of nucleotide and indicate a mixture of NTD 'up' and 'down' states. The NMR data thus indicate that the D1 binding pocket is still able to interact with nucleotide but not to structurally discriminate ATP($\gamma$S) from ADP. The binding affinity of K251A mutant for ADP is below the detection limit of ITC (Table S7).

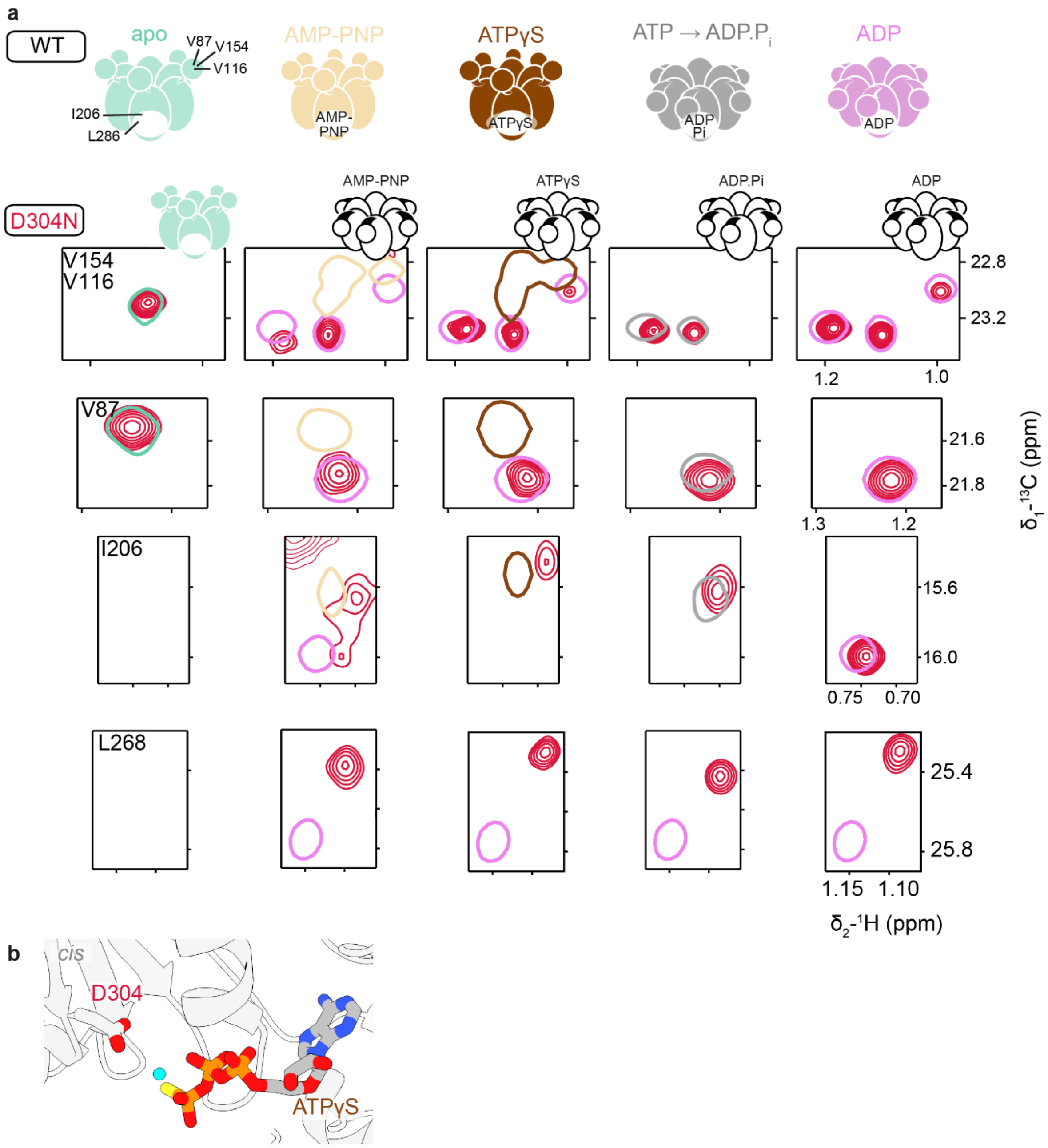

**Supplementary Fig. 6 NMR spectra of p97-ND1L D304N. a**, Selected spectral regions from ($^1$H,$^{13}$C)-HMQC spectra of p97-ND1L D304N as a function of nucleotide present. Residues V116, V154 and V87 report on the NTD position, while residues I206 and L268 reflect the conformation of the D1 binding pocket. Methyl group correlations from the respective wt spectra are indicated in single contours; when wt and mutant spectra differ, the corresponding wt-ADP spectra are shown in addition. **b**, Close-up of the D1 binding pocket in ATPγS state. D304 is crucial for $Mg^{2+}$ binding. The D304N mutant binds nucleotide (*cf*. Table S7 and NMR signals of I206/L268 in nucleotide presence *vs*. absence), yet it fails to assume an NTD 'up' state in the presence of ATPγS/AMP-PNP. The coordination of $Mg^{2+}$ is a prerequisite to structurally recognize ATP and its analogues. Due to its low ATPase rate (Fig. 3b), it remains unclear whether the D304N mutant is able to hydrolyse ATP at all.

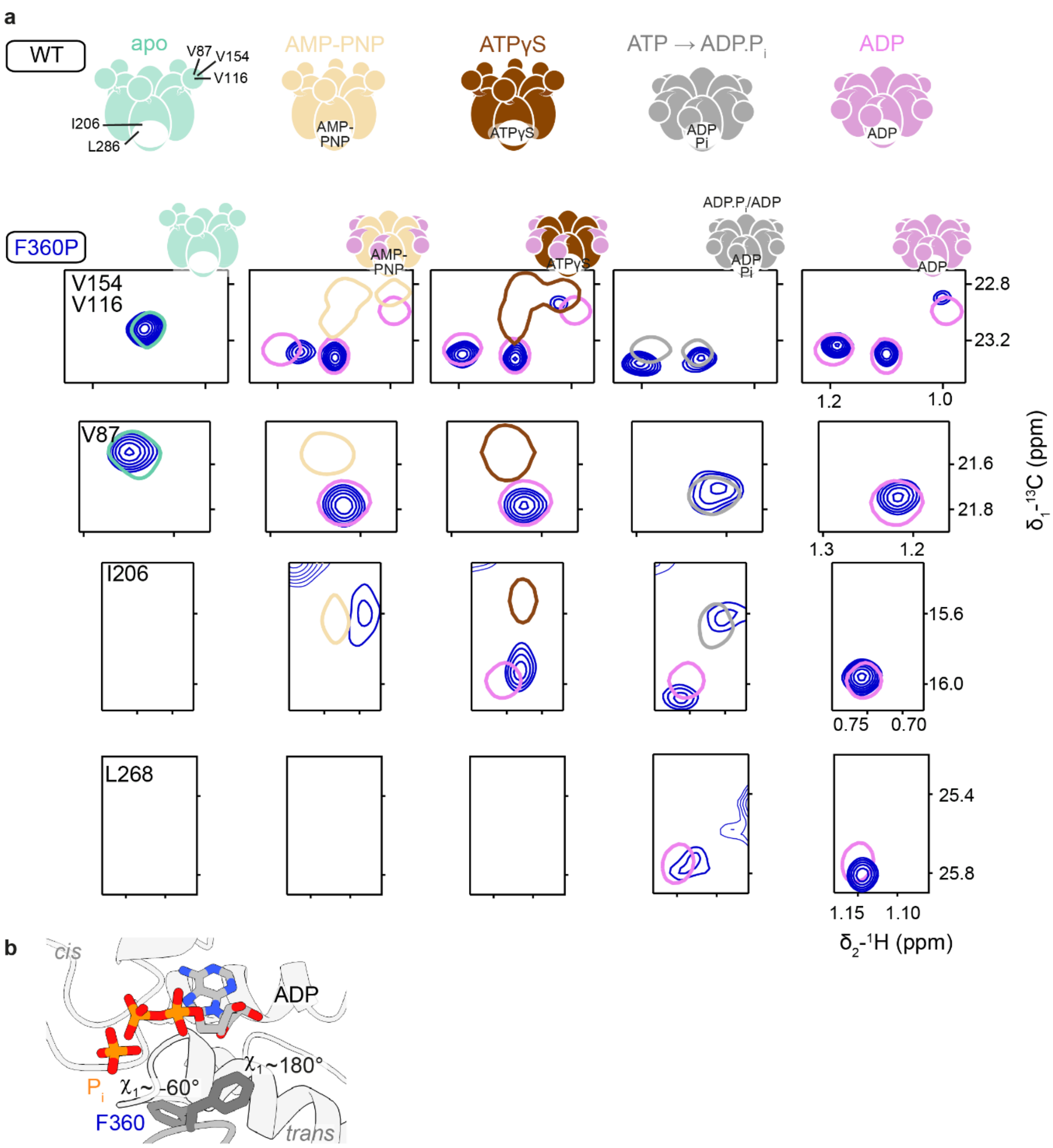

**Supplementary Fig. 7 NMR spectra of p97-ND1L F360P.** **a**, Selected spectral regions from ($^{1}H$,$^{13}C$)-HMQC spectra of p97-ND1L F360P as a function of nucleotide present. Residues V116, V154 and V87 report on the NTD position ('up' *vs.* 'down'), while residues I206 and L268 reflect the conformation of the D1 binding pocket and its nucleotide. Methyl group correlations from the respective wt spectra are indicated in single contours; when wt and mutant spectra differ, the corresponding wt-ADP spectra are shown in addition. **b**, Close-up of the D1 binding pocket in ADP.$P_i$ state. F360 switches between different rotamers: $\chi_1$~180° allows for association with the neighbouring helix $\alpha_{407-423}$, while $\chi_1$~-60° leads to disassociation. The F360P mutation decouples the NTD position from the D1 nucleotide state, evidenced by the NTD 'down' position in the presence of AMP-PNP and ATP$\gamma$S. Spectra recorded in the presence of ATP show a mixture of pre-hydrolysis (AMP-PNP-like) and post-hydrolysis (ADP-like) states for residue I206.

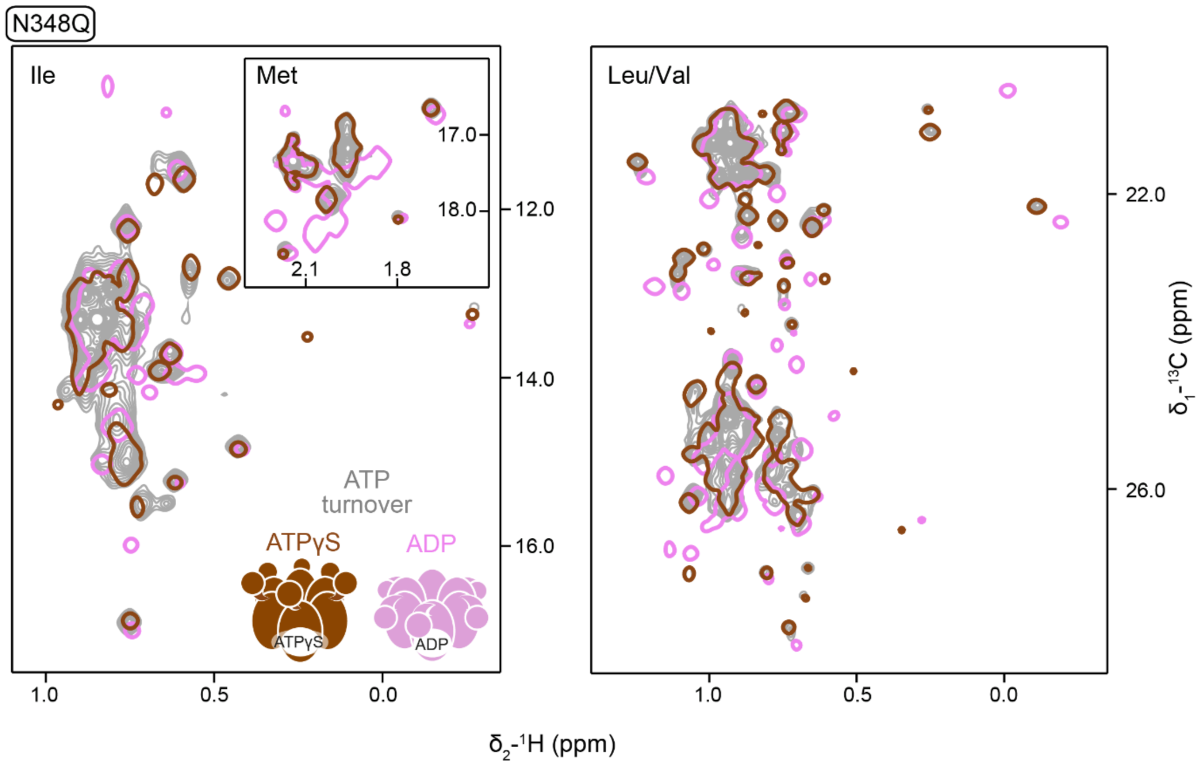

**Supplementary Fig. 8 NMR spectra of p97-ND1L N348Q.** ($^{1}H$,$^{13}C$)-HMQC spectra of *proR*-$^{13}CH_3$-ILVM-labelled p97-ND1L N348Q. The spectrum recorded in the presence of the ATP-regeneration system (multiple grey contours) is very similar to the spectrum recorded in the presence of slowly-hydrolysable ATPγS (single brown contours), documenting the inability of the N348Q mutant to hydrolyse ATP.

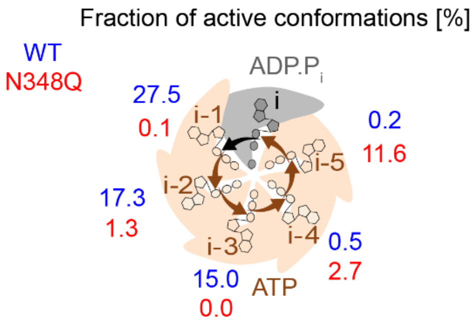

**Supplementary Fig. 9 Fraction of reaction-competent conformations.** Conformations poised for ATP hydrolysis as defined in Fig. 3e were identified at each ATP-bound D1 active site over all frames of the full MD simulation of the wt protein and the N348Q mutant. Reaction-competent conformations are found in a significantly higher number of frames for the wt protein than for the mutant. The single ADP.$P_i$-bound subunit breaks the symmetry of the hexamer. In the simulation of the wt, the fraction of reactive conformations in each subunit declines from the ADP.$P_i$-bound subunit in a counter clockwise manner. This distribution is reminiscent of the unidirectional hydrolysis mechanism postulated for many AAA proteins based on cryo-EM structures[39]. However, testing the significance of this observation will require more extensive statistics.

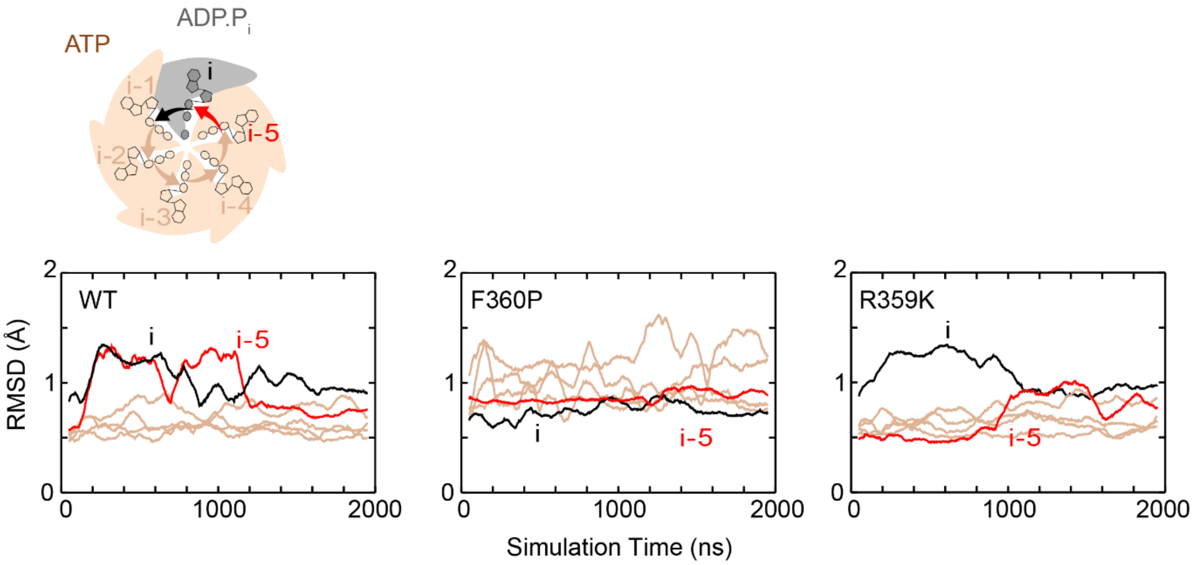

**Supplementary Fig. 10 Conformational changes in the sensor loop.** RMSD (root-mean-quare deviation) fluctuations indicate the deviation of the backbone of sensor loop residues (348-361) from their position in the first frame over the course of the 2 μs-MD trajectory. In the wt protein, the loop displays increased mobility if one of the two adjacent active sites is occupied by ADP.P$_i$ (red/black) compared to ATP (brown). In the hyperactive F360P mutant, the mobility of the loop is overall increased, irrespective of nucleotide state. In the inactive R359K mutants, loop mobility is overall decreased.

**a**

| Mutant | wt | P246T | P247A | P247K | K251A | D304N | E305Q | N348Q | R349A | R359A | R359K | F360A | F360P | R362A |
|---|---|---|---|---|---|---|---|---|---|---|---|---|---|---|
| NTD down motion upon $P_i$ binding | v | x | v | ~ | ~ | v | ~ | v | v | ~ | v | v | v | x |
| Hexamer in apo state | v | v | v | x | v | v | ~ | v | v | x | v | v | v | x |

| | |
|---|---|
| v | yes |
| x | no |
| ~ | partial |

**b**

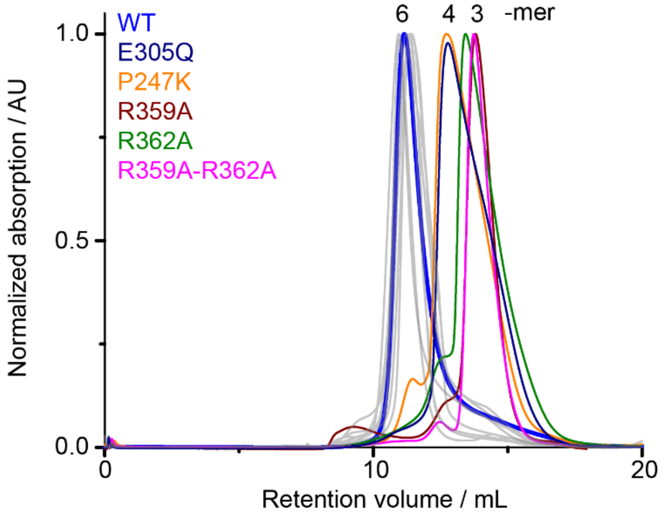

**Supplementary Fig. 11 Comparison of point mutants in apo state. a**, Structural response of p97-ND1L mutants to $P_i$ ion addition and oligomeric states of the respective mutants in the absence of nucleotide. The response of a given mutant to $P_i$ binding is linked to its ability to form intact hexamers in apo state. **b**, The oligomeric states of p97-ND1L mutants in the absence of nucleotide were assessed by SEC. Non-hexameric mutants and the wt are highlighted in colour, all others are shown in grey. P274K and R359A require nucleotide for hexamerisation, R362A mutants do not form hexamers at all (Fig. E4).

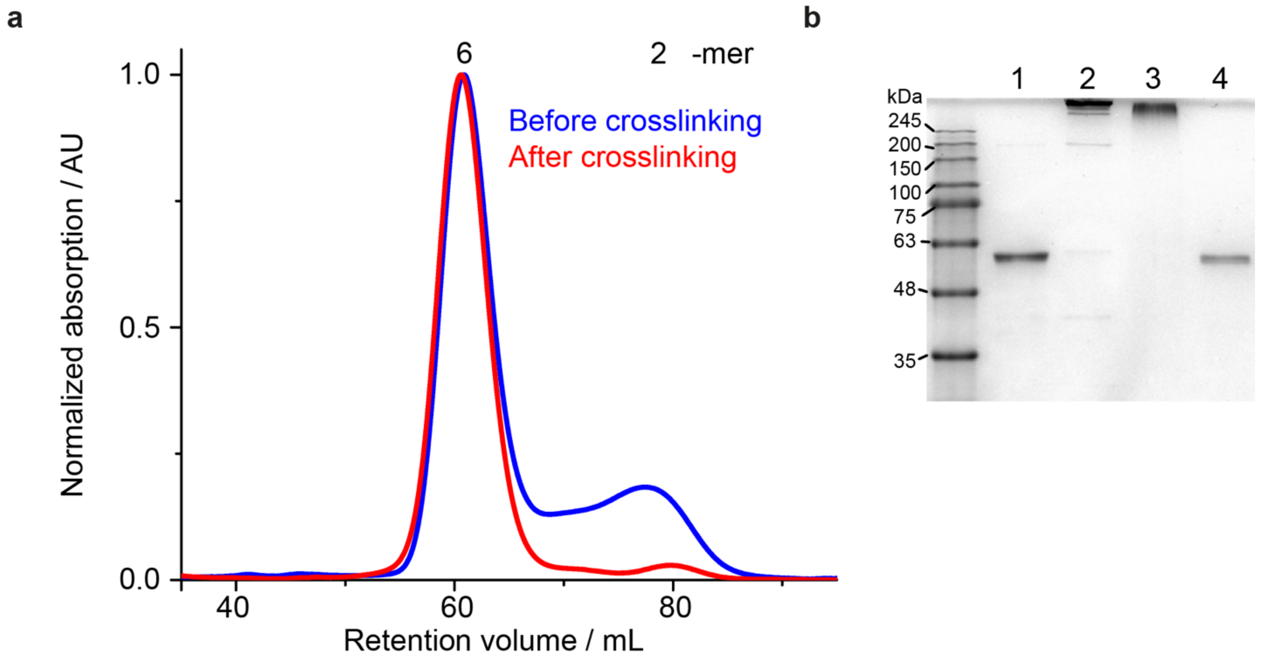

**Supplementary Fig. 12 Crosslinking of the ∆Cys-F360C-A413C mutant.**

**a**, Oligomeric state of the p97-ND1L ∆Cys-F360C-A413C mutant assessed by SEC before and after crosslinking with BMOE. In both cases the protein elutes at a volume corresponding to a hexameric assembly. **b**, SDS-PAGE confirming the crosslinking. 1: peak from SEC corresponding to hexameric p97 before crosslinking; 2: crosslinking reaction mixture; 3: peak from SEC corresponding to hexameric p97 after crosslinking; 4: ND1L wt reference.

### Supplementary Tables

**Supplementary Table 1.** Experimental setup for solution-state NMR data acquisition.

| | |
|---|---|
| Spectrometer $^{1}H$ frequency [MHz] | 800, 900, 950 |
| Probe | cryo-TCI |
| Pulse sequence | ($^{1}H$,$^{13}C$)-HMQC |
| Number of scans | 4 |
| Acquisition time [min] | 42 |
| Spectral width [ppm] | 16.0($^{1}H$) / 25.0($^{13}C$) |
| Transmitter frequency offset [ppm] | 4.707($^{1}H$) / 18.6($^{13}C$) |

**Supplementary Table 2.** Experimental setup for solid-state NMR data acquisition.

| | | |
|---|---|---|
| Spectrometer $^{1}H$ frequency [MHz] | 800 | |
| Probe | Bruker 1.3 mm WB in HPC mode | |
| MAS rate | 45 kHz | |
| Pulse sequence | Direct excitation | Cross-polarization |
| Description | $^{31}P$ direct excitation spectra, $^{1}H$ decoupling during acquisition | $^{1}H$-$^{31}P$ cross-polarization spectra, $^{1}H$ decoupling during acquisition |
| 90° excitation pulse [µs] | 2.0 ($^{31}P$) | 1.59 ($^{1}H$) |
| Cross-polarization transfer | | Ramp 70-100%, 30 kHz ($^{31}P$), 93 kHz ($^{1}H$) |
| Transmitter frequency offset [ppm] | 0 ($^{31}P$), 4.0 ($^{1}H$) | 0 (ATP) / 15 (ATPγS) ($^{31}P$), 4.0 ($^{1}H$) |
| $^{1}H$ decoupling during acquisition | sITPPM, 11 kHz | |
| Acquisition time [ms] | 41 ($^{31}P$) | 20.5 ($^{31}P$) |
| Number of scans/ experimental time [min] | 512/69 | 8192/140 |
| Spectral width [ppm] | 154.3 | |
| Interscan delay [s] | 8 | 1 |

**Supplementary Table 3.** Cryo-EM data collection and processing statistics.

| **Sample** | fl p97 in ATP-regenerating buffer system | |
|---|---|---|
| **Data Collection** | | |
| Microscope | Titan Krios | |
| Voltage (kV) | 300 | |
| Magnification | 165,000 | |
| Pixel size (Å) | 0.72 | |
| Exposure rate ( e/Å$^2$/s) | 10.4 | |
| Total Exposure (e/Å$^2$) | 40 | |
| Nominal defocus range (μm) | -0.9 to -2.2 | |
| Energy filter and detector | Selectris and Falcon 4 | |
| Energy filter slit (eV) | 15 | |
| Number of images | 10011 | |
| Number of EER frames | 931 | |
| **Image Processing** | | |
| Name of the maps | D1-D2 Focused | NTD Focused |
| Symmetry imposed | C6 | C1 |
| Initial particle number | 693,686 | 693,686 |
| Final particle number | 86,760 | 112,231 |
| FSC threshold | 0.143 | 0.143 |
| Map resolution (Å) | 2.61 | 3.27 |
| Map resolution range (Å) | 2.53 – 4.03 | 3.14 - 5.41 |
| Map sharpening B factor (Å$^2$) | -58 | -67 |
| EMDB entry | EMD-16782 | EMD-16781 |

**Supplementary Table 4.** Cryo-EM model building.

| **Model statistics** | |
|---|---|
| Initial model used (PDB code) | 5ftl, 5ftm[10] |
| Nonhydrogen atoms | 35334 |
| Chains | 6 |
| Protein residues | 4404 |
| Water | 246 |
| Ligands | $K^+$: 6, $PO_4^{3-}$: 6, $Mg^{2+}$: 12, ATP: 6, ADP: 6 |
| R.m.s deviations<br>Bond length (Å)<br>Bond angles (°) | <br>0.0041<br>0.72 |
| Ramachandran plot<br>Favored (%)<br>Allowed (%)<br>Outliers (%) | <br>95.19<br>4.53<br>0.27 |
| Rotamer outliers (%) | 0 |
| Clashscore | 18.17 |
| Molprobity score[40] | 2.09 |
| PDB entry | 8cpf |

**Supplementary Table 5.** Overview of all MD simulations. All simulations were performed on p97 hexamers.

| Simulation Name | System | Template | Sampling time |
|---|---|---|---|
| **ADP.$P_i$ prediction** | ND1L, 5xATP, 1xADP.$P_i$ ($HPO_4^{2-}$) | 4ko8 | 2000 ns |
| **ADP.$P_i$ prediction** | ND1L, 5xATP, 1xADP.$P_i$ ($H_2PO_4^-$) | 4ko8 | 1000 ns |
| **R359K mutant** | ND1L, 5xATP, 1xADP.$P_i$, R359K | 4ko8 | 2000 ns |
| **F360P mutant** | ND1L, 5xATP, 1xADP.$P_i$, F360P | 4ko8 | 2000 ns |
| **N348Q mutant** | ND1L, 5xATP, 1xADP.$P_i$, N448Q | 4ko8 | 2000 ns |
| **$P_i$ + $P_i$** | D1, 5xATP, 1x$P_i$+$P_i$, no $Mg^{2+}$ | 4ko8 | 1000 ns |
| **ADP.$P_i$ refinement** | full-length, 6xADP.$P_i$ (D1), 6xATP (D2) | This work | 1300 ns |
| **$P_i$ dissociation** | Like above, no $Mg^{2+}$ | This work | 700 ns |

**Supplementary Table 6.** Overview of the seven equilibration steps performed for all simulations. The initial minimization is followed by six equilibration simulations with decreasing force constants (K) on positional restraints of amino acid residues, ATP, ADP, $P_i$ and $Mg^{2+}$ ions.

| | Steps | Time step | K solute | Temp. | Pressure |
|---|---|---|---|---|---|
| **min** | 1000 | minimization | 10.0 | --- | --- |
| **Eq1** | 25000 | 1fs | 10.0 | 303.15 K | NA |
| **Eq2** | 25000 | 1fs | 5.0 | 303.15 K | NA |
| **Eq3** | 25000 | 1fs | 2.5 | 303.15 K | 1 atm |
| **Eq4** | 50000 | 2fs | 1.0 | 303.15 K | 1 atm |
| **Eq5** | 50000 | 2fs | 0.5 | 303.15 K | 1 atm |
| **Eq6** | 250000 | 2fs | 0.1 | 303.15 K | 1 atm |

**Supplementary Table 7.** Binding affinities of p97-ND1L wt and mutants towards ADP and ATPγS determined by ITC. Abbreviations: n.d. no binding detected; n number of replicates. In the absence of $Mg^{2+}$ ions, p97 cannot bind ATPγS with high affinity. This experiment serves as a negative control.

| p97-ND1L | ADP $K_D$ (μM) | ATPγS $K_D$ (μM) |
|---|---|---|
| WT | 0.08 ± 0.01 (n=3) | 0.19±0.07 (n=3) |
| P246T | 0.02 ± 0.0007 (n=2) | 0.09 ± 0.05 (n=3) |
| P247A | 0.08 ± 0.02 (n=3) | 0.18 ± 0.16 (n=2) |
| P247K | 0.05 ± 0.03 (n=3) | 0.01 ± 0.004 (n=2) |
| D304N | 0.05 ± 0.002 (n=2) | 0.30 ± 0.08 (n=3) |
| E305Q | 0.32 ± 0.001 (n=2) | 0.08 ± 0.001 (n=3) |
| N348Q | 0.18 ± 0.04 (n=3) | 0.68 ± 0.06 (n=2) |
| R359A | 1.35 ± 0.78 (n=3) | 0.04 ± 0.02 (n=3) |
| R359K | 0.11 ± 0.03 (n=3) | 0.21 ± 0.16 (n=3) |
| F360A | 0.06 ± 0.03 (n=3) | 0.23 ± 0.01 (n=2) |
| F360P | 0.17 ± 0.02 (n=3) | 0.14 ± 0.02 (n=2) |
| R359A-R362A | 0.05 ± 0.03 (n=3) | 0.11 ± 0.02 (n=2) |
| K251A | n.d. (n=2) | |
| wt without $Mg^{2+}$ | | 1.08 ± 0.007 (n=2) |

### Supplementary Videos

**Supplementary Video 1.** Transition from ADP.$P_i$ state A to state B observed in MD simulations.

Close up of the D1 active site from an MD simulation sampled immediately after *in silico*-transformation of ATP into ADP and $P_i$ for 2 µs. A concerted rotamer flip of residues R359 and F360 (bottom) is observed after ~1.2 µs, marking the transition from ADP.$P_i$ state A to state B. This switch is accompanied by a shift in the position of the cleaved $P_i$ ion and of the $Mg^{2+}$ ion (pink), which stably bridges the $P_i$ and ADP (top).

**Supplementary Video 2.** The R359K mutant fails to transition to state B in MD simulations.

Close up of the D1 active site bearing the R359K mutation from an MD simulation sampled immediately after *in silico*-transformation of ATP into ADP and $P_i$ for 2 µs. K359 is positioned in between ADP and the cleaved $P_i$ ion, symmetrically opposite to K251 (*c.f.* Fig. E8b). In contrast to the wt protein, only a single ADP.$P_i$ geometry is observed, which is very stable (*c.f.* Fig 5a) and similar to state A with respect to the positioning of the $Mg^{2+}$ and $P_i$ ions and the dissociation of F360 from helix $\alpha_{407\text{-}423}$. The R359K mutant does not release the reaction products efficiently.

**Supplementary Video 3.** The F360P mutant transitions from state A to state B in MD simulations.

Close up of the D1 active site with F360P mutation from an MD simulation sampled immediately *in silico*-transformation of ATP into ADP and $P_i$ for 2 µs. The $P_i$ and $Mg^{2+}$ ions move between positions that are similar to ADP.$P_i$ states A and B of the wt. However, the absence of the F360 side chain prevents the coupling of ATP-hydrolysis events to dissociation from helix $\alpha_{407\text{-}423}$.

**Supplementary Video 4.** ADP.$P_i$ state can be mimicked by two $P_i$ ions.

Close up of the D1 nucleotide binding pocket in apo state in the presence of exogeneous $P_i$ ions. The ADP.$P_i$ state is mimicked by two $P_i$ ions bridged by $K^+$ ions from the solvent (green), which exchange frequently during the 1 µs simulation time. The $P_i$ ions occupy the same positions as the β-P of ADP and the cleaved $P_i$ ion. A $Mg^{2+}$ ion is not required for stabilization of the arrangement.

**Supplementary Video 5.** Long-range structural transitions associated with ATP hydrolysis in D1.

The NTD moves upon ATP hydrolysis in D1, starting from the 'up 'position observed in the ATPγS state (PDB: 5ftn[10]) to the 'down' position in ADP.$P_i$ states A and B (PDB: 8cpf, this work). The active site is located at the inter-subunit interface, with the *cis*-acting subunit shown in purple and the *trans*-acting in pink. The transition between ADP.$P_i$ states A and B is evident in rotamer switches of R359 and F360 and repositioning of the $P_i$ and $Mg^{2+}$ ions. Free energy calculations (c.f. Fig. 5a) suggest that state B marks the onset to dissociation. H384 is located next to the ribose moiety of the nucleotide and displays two side chain rotamers in the ADP.$P_i$ state, which cannot be assigned to states A or B with certainty. In the NTD 'down' position, electrostatic interactions, which form only after ATP hydrolysis, stabilize the NTD-D1 interface: between E34 and K386 as well as between R155 and N387.

### Supplementary References

1. Rydzek S, Shein M, Bielytskyi P, Schutz AK. Observation of a Transient Reaction Intermediate Illuminates the Mechanochemical Cycle of the AAA-ATPase p97. *J Am Chem Soc* 2020, **142**(34): 14472-14480.

2. Schuetz AK, Kay LE. A Dynamic molecular basis for malfunction in disease mutants of p97/VCP. *Elife* 2016, **5**.

3. Jensen LM, Walker EJ, Ghildyal R (eds). *Rhinoviruses, Methods and Protocols*, 2014.

4. Vranken WF, Boucher W, Stevens TJ, Fogh RH, Pajon A, Llinas M*, et al.* The CCPN data model for NMR spectroscopy: development of a software pipeline. *Proteins* 2005, **59**(4): 687-696.

5. Zheng SQ, Palovcak E, Armache JP, Verba KA, Cheng Y, Agard DA. MotionCor2: anisotropic correction of beam-induced motion for improved cryo-electron microscopy. *Nat Methods* 2017, **14**(4): 331-332.

6. Zivanov J, Nakane T, Forsberg BO, Kimanius D, Hagen WJ, Lindahl E*, et al.* New tools for automated high-resolution cryo-EM structure determination in RELION-3. *Elife* 2018, **7**: e42166.

7. Rohou A, Grigorieff N. CTFFIND4: Fast and accurate defocus estimation from electron micrographs. *J Struct Biol* 2015, **192**(2): 216-221.

8. Wagner T, Merino F, Stabrin M, Moriya T, Antoni C, Apelbaum A*, et al.* SPHIRE-crYOLO is a fast and accurate fully automated particle picker for cryo-EM. *Commun Biology* 2019, **2**(1): 218.

9. Kimanius D, Dong L, Sharov G, Nakane T, Scheres SHW. New tools for automated cryo-EM single-particle analysis in RELION-4.0. *Biochem J* 2021, **478**(24): 4169-4185.

10. Banerjee S, Bartesaghi A, Merk A, Rao P, Bulfer SL, Yan Y*, et al.* 2.3 A resolution cryo-EM structure of human p97 and mechanism of allosteric inhibition. *Science* 2016, **351**(6275): 871-875.

11. Emsley P, Lohkamp B, Scott WG, Cowtan K. Features and development of Coot. *Acta Crystallogr D Biol Crystallogr* 2010, **66**(Pt 4): 486-501.

12. Liebschner D, Afonine PV, Baker ML, Bunkoczi G, Chen VB, Croll TI*, et al.* Macromolecular structure determination using X-rays, neutrons and electrons: recent developments in Phenix. *Acta Crystallogr D Struct Biol* 2019, **75**(Pt 10): 861-877.

13. Salomon-Ferrer R, Gotz AW, Poole D, Le Grand S, Walker RC. Routine Microsecond Molecular Dynamics Simulations with AMBER on GPUs. 2. Explicit Solvent Particle Mesh Ewald. *J Chem Theory Comput* 2013, **9**(9): 3878-3888.

14. AMBER 2018. University of California, San Francisco; 2018.

15. Maier JA, Martinez C, Kasavajhala K, Wickstrom L, Hauser KE, Simmerling C. ff14SB: Improving the Accuracy of Protein Side Chain and Backbone Parameters from ff99SB. *J Chem Theory Comput* 2015, **11**(8): 3696-3713.

16. Berendsen HJC, Grigera JR, Straatsma TP. The missing term in effective pair potentials. *J Phys Chem* 1987, **91**(24): 6269-6271.

17. Li P, Song LF, Merz KM, Jr. Parameterization of highly charged metal ions using the 12-6-4 LJ-type nonbonded model in explicit water. *J Phys Chem B* 2015, **119**(3): 883-895.

18. Meagher KL, Redman LT, Carlson HA. Development of polyphosphate parameters for use with the AMBER force field. *J Comput Chem* 2003, **24**(9)**:** 1016-1025.

19. Wang J, Wolf RM, Caldwell JW, Kollman PA, Case DA. Development and testing of a general amber force field. *J Comput Chem* 2004, **25**(9)**:** 1157-1174.

20. Fox T, Kollman PA. Application of the RESP Methodology in the Parametrization of Organic Solvents. *J Phys Chem B* 1998, **102**(41)**:** 8070-8079.

21. Gaussian 09, Revision A.02. Wallingford CT; 2016.

22. Goga N, Rzepiela AJ, de Vries AH, Marrink SJ, Berendsen HJ. Efficient Algorithms for Langevin and DPD Dynamics. *J Chem Theory Comput* 2012, **8**(10)**:** 3637-3649.

23. Åqvist J, Wennerström P, Nervall M, Bjelic S, Brandsdal BO. Molecular dynamics simulations of water and biomolecules with a Monte Carlo constant pressure algorithm. *Chem Phys Lett* 2004, **384**(4-6)**:** 288-294.

24. Darden T, York D, Pedersen L. Particle mesh Ewald: AnN·log(N) method for Ewald sums in large systems. *J Chem Phys* 1993, **98**(12)**:** 10089-10092.

25. Andersen HC. Rattle: A “velocity” version of the shake algorithm for molecular dynamics calculations. *J Comput Phys* 1983, **52**(1)**:** 24-34.

26. Hopkins CW, Grand SL, Walker RC, Roitberg AE. Long-Time-Step Molecular Dynamics through Hydrogen Mass Repartitioning. *Journal of Chemical Theory and Computation* 2015, **11**(4)**:** 1864-1874.

27. Humphrey W, Dalke A, Schulten K. VMD: visual molecular dynamics. *J Mol Graph* 1996, **14**(1)**:** 33-38, 27-38.

28. Roe DR, Cheatham TE, 3rd. PTRAJ and CPPTRAJ: Software for Processing and Analysis of Molecular Dynamics Trajectory Data. *J Chem Theory Comput* 2013, **9**(7)**:** 3084-3095.

29. Tang WK, Xia D. Altered intersubunit communication is the molecular basis for functional defects of pathogenic p97 mutants. *J Biol Chem* 2013, **288**(51)**:** 36624-36635.

30. Jurrus E, Engel D, Star K, Monson K, Brandi J, Felberg LE*, et al.* Improvements to the APBS biomolecular solvation software suite. *Protein Sci* 2018, **27**(1)**:** 112-128.

31. Miller BR, 3rd, McGee TD, Jr., Swails JM, Homeyer N, Gohlke H, Roitberg AE. MMPBSA.py: An Efficient Program for End-State Free Energy Calculations. *J Chem Theory Comput* 2012, **8**(9)**:** 3314-3321.

32. Dolinsky TJ, Czodrowski P, Li H, Nielsen JE, Jensen JH, Klebe G*, et al.* PDB2PQR: expanding and upgrading automated preparation of biomolecular structures for molecular simulations. *Nucleic Acids Res* 2007, **35**(Web Server issue)**:** W522-525.

33. Sievers F, Wilm A, Dineen D, Gibson TJ, Karplus K, Li W*, et al.* Fast, scalable generation of high-quality protein multiple sequence alignments using Clustal Omega. *Mol Syst Biol* 2011, **7**(1)**:** 539.

34. Waterhouse AM, Procter JB, Martin DM, Clamp M, Barton GJ. Jalview Version 2--a multiple sequence alignment editor and analysis workbench. *Bioinformatics* 2009, **25**(9)**:** 1189-1191.

35. Pettersen EF, Goddard TD, Huang CC, Couch GS, Greenblatt DM, Meng EC*, et al.* UCSF Chimera--a visualization system for exploratory research and analysis. *J Comput Chem* 2004, **25**(13)**:** 1605-1612.

36. Pettersen EF, Goddard TD, Huang CC, Meng EC, Couch GS, Croll TI*, et al.* UCSF ChimeraX: Structure visualization for researchers, educators, and developers. *Protein Science* 2021, **30**(1)**:** 70-82.

37. Chen S, McMullan G, Faruqi AR, Murshudov GN, Short JM, Scheres SH*, et al.* High-resolution noise substitution to measure overfitting and validate resolution in 3D structure determination by single particle electron cryomicroscopy. *Ultramicroscopy* 2013, **135**(C)**:** 24-35.

38. Wendler P, Ciniawsky S, Kock M, Kube S. Structure and function of the AAA+ nucleotide binding pocket. *Biochim Biophys Acta* 2012, **1823**(1)**:** 2-14.

39. Seraphim TV, Houry WA. AAA+ proteins. *Curr Biol* 2020, **30**(6)**:** R251-R257.

40. Davis IW, Leaver-Fay A, Chen VB, Block JN, Kapral GJ, Wang X*, et al.* MolProbity: all-atom contacts and structure validation for proteins and nucleic acids. *Nucleic Acids Res* 2007, **35**(Web Server issue)**:** W375-383.
